# Effective High-Accuracy Prediction of Protein Structures from Easily Obtainable Artificial Homologous Sequences by Structure-Stability-Based Selection

**DOI:** 10.1101/2023.11.22.568372

**Authors:** Jinle Tang, Zhe Zhang, Jian Zhan, Yaoqi Zhou

## Abstract

High-resolution protein structure determination by experimental techniques is notoriously costly and labor intensive. This problem is mostly solved with arrival of deep-learning-based computational prediction by AlphaFold2 but only for those proteins with enough naturally occurring homologous sequences. Here, we attempt to close the remaining gap by employing artificially generated, structure-stability-selected homologous sequences as an input for AlphaFold2. We showed that only one round of selection of deeply mutated sequences of a few mutations is sufficient to bring the accuracy of predicted structures to better than 2 Å RMSD from their respective native structures for four of the five proteins experimented. The performance for three out of five proteins is even better than AlphaFold2 with naturally occurring sequences. The only protein with predicted structure of >2 Å (at 2.92 Å) RMSD is due to a fully exposed (i.e., likely flexible) β-hairpin. The result supports a future of determining protein structures at low cost and fast turnaround by integrating simple molecular biology experiments (deep mutational scanning and *in vivo* or *in vitro* selection) with high-throughput sequencing. The technique proposed here can be further extended to predict structures of protein complexes as well as proteins with posttranslational modifications.

## Introduction

Recently, AlphaFold2 achieved something thought impossible: the accuracy of predicted protein structures matches that of experimental structure determination (1). This unexpected high-accuracy performance was achieved by end-to-end large-scale learning for mapping between multiple homologous sequences and the corresponding “single” structure (2). That is, AlphaFold2 relies strongly on evolutionary and co-evolutionary signals encoded in the multiple sequence alignments (MSAs). A large reduction in accuracy was observed in AlphaFold2 prediction for those proteins with a median alignment depth of < 30 sequences, measured by removing highly homologous sequences using a cutoff of 80% sequence identity (1). Many proteins or protein regions, newly evolved proteins, orphan proteins and antibodies, in particular, do not have that many homologous sequences. Although it was estimated that AlphaFold2 can cover about 98.5% of the human proteome, only 58% residues could be predicted with confidence, while mere 36% residues could be predicted with high confidence (3, 4).

One way to address the proteins with insufficient homologous sequences is to update the sequence database with newly solved genomic or meta-genomic data and to increase homolog-searching sensitivity (5–7). This may work for some proteins but does not solve the problem for those proteins lacking naturally occurring homologous sequences. Another approach is to employ protein language models containing evolution information implicitly by learning from other evolved sequences in the database (8–11). These methods, however, do not yet have a consistent, high-accuracy prediction for different proteins.

If natural homologous sequences are not sufficient, can we employ homologous sequences generated artificially for improving structure prediction? Unlike natural sequences, most of which evolved over billions of years, artificially generated homologous sequences were resulted from a short duration of laboratory evolution with a few mutations (typically having >95% sequence identities) and thus require many more mutation variants than homologs with <85% sequence identities to cover key correlated mutations between different sequence positions. Thanks to high-throughput sequencing, deep mutational scanning became a popular tool for the efficient assessment of protein function and generating a large number of artificial homologs from experiments (12, 13). These homologs with a few mutations have been demonstrated useful in epistasis (14, 15) or covariation-induced deviation (16) analysis and generation of distance restraints for improved protein and RNA structure prediction (14–18). It is not yet clear, however, if these highly homologous sequences (>95% sequence identity) can be directly employed in AlphaFold2 or related predictors that were trained mostly by naturally occurring homologs with low sequence identity (<85% sequence identity) to a query sequence and between each other.

Moreover, most deep mutational scanning experiments were conducted by functional selections such as antibiotic resistance proteins (17, 18), fluorescence proteins (19, 20), and proteins with binding properties (21, 22). Given thousands of protein functions, it is not feasible to develop different selection techniques for different functions, not to mention that not all protein functions are suitable for high-throughput studies. Thus, harnessing artificial homologous sequences for structural inference would call for a technique that selects those structurally homologous sequences based on structural stability. Such a technique in principle would work for all proteins that rely on stable structures for function, regardless the type of function. Existing methods that monitor the folding stability of proteins rely on the utilization of proteases induced proteolysis (23, 24) or protein-fragment complementation assay (25–28). However, each technique has its limitation such as the capacity of selection or the size limitation of target proteins. More importantly, these techniques were not designed or tested for protein structure prediction.

Here, we developed an efficient, cost-effective assay for Structural Inference By Structure-stability-based selection of deep mutation variants and high throughput sequencing (Sibs-Seq). We demonstrated that unlike naturally occurring homologous sequences evolved over billions of years, these stability-selected, artificial homologs in a single-round 24 or 36 hours of mutation selections is often sufficient for producing high-accuracy structure prediction with < 2 Å root-mean-squared deviation (RMSD) to the native structure.

## Results

### The flowchart of Sibs-Seq

Figure 1a demonstrates the overall workflow of Sibs-Seq for protein structural interference. A target protein of interest (POI) is subjected to error prone PCR (EP-PCR) to construct a mutation library (>10^6^) that is inserted between two assisted-complementary fragments of the murine dihydrofolate reductase (mDHFR: N terminal fragment, (mF1, 2) and the C terminal fragment (mF3)), which can be reconstituted into a functional mDHFR when they are brought into proximity by a stably folded POI (29–32). A functional mDHFR will then produce tetrahydrofolate (THF) essential for Trimethoprim (TMP) resistance and allow growth of TMP-sensitive *E. coli* in the presence of TMP (29, 33). By contrast, if the POI is in an unstable structural state, two fragments of mDHFR would be too far apart to reconstitute into a functional mDHFR. Lacking a functional mDHFR will inhibit the growth of TMP-sensitive *E. coli* cells when treated with TMP. Thus, when inserting a library of randomly mutated POIs, those stably folded variants will lead to TMP-resistant *E. coli* that can grow in the presence of TMP. The sequence counts of each mutant available from high-throughput sequencing will allow to calculate its fitness score, which provides an estimation for its stability. Then, these artificially generated, structurally stable variants can be aligned as an input (artificial MSA, aMSA) for structure prediction by AlphaFold2.

**Figure 1.**
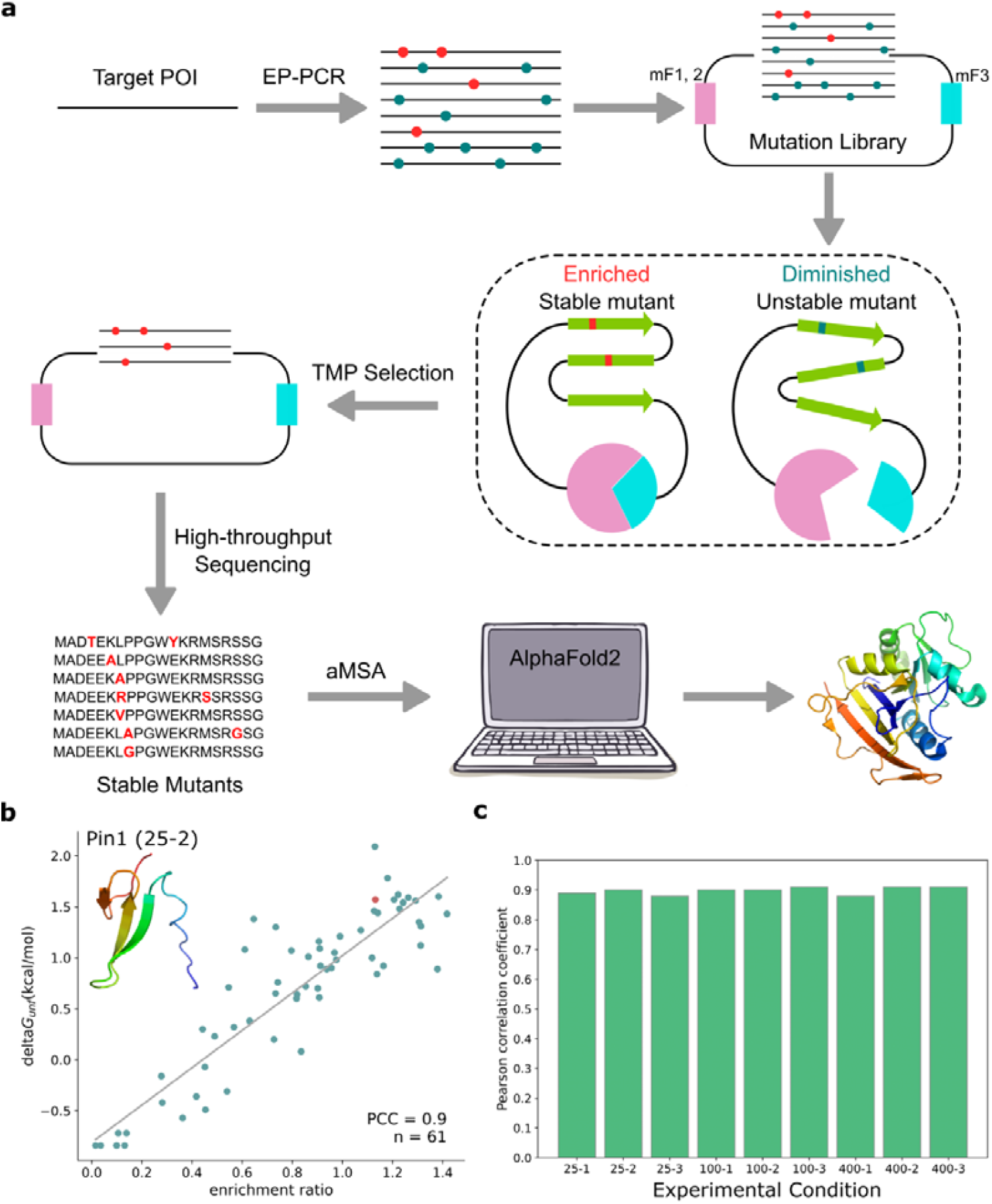
Design and validation of Sibs-Seq for structure inference by stability-based selection and enrichment of stable mutants of a randomly mutated target protein of interest (POI). **a**, A target protein of interest is mutated by error-prone PCR and inserted into two assisted-complementary split position of a murine dihydrofolate reductase (mF1,2 and mF3 of mDHFR). These two fragments will reconstitute for structurally stable variants of POIs into a functional mDHFR and lead to TMP resistance *E. coli*, whereas an unstable POI variant is unable to bring two fragments of mDHFR together for a functional mDHFR and lead to TMP sensitive *E. coli*. TMP-selected stable variants can be revealed by high-throughput sequencing and employed as artificial homologous sequences (aMSA) for structure prediction by AlphaFold2. **b, c**, The Sibs-Seq assay was validated by using 61 variants of Pin1 WW domain with known stabilities (ΔG_un_ in kcal/mol, measured by thermal denaturation). **b** shows the relation between ΔG_un_ of a mutant and its sequence enrichment ratio at the experimental condition of 25 μg/ml TMP for 2 × 12 h with Pearson Correlation Coefficient, PCC = 0.9 for Pin1 WW domain. The structure was drawn according to PDB ID: 2M8I for Pin1 WW domain. The red dot represents the position of the wild-type sequence. **c** displays high PCC values for all selection pressures tested (TMP concentrations from 25 μg/ml, 100 μg/ml, to 400 μg/ml and incubation time from 1 × 12 h, 2 × 12 h, to 3 × 12 h).

### Validation on enrichment of structural stable variants by Sibs-Seq

To validate Sibs-Seq, we purchased a synthetic DNA library encoding 61 variants of 50-residue Pin1 WW-domain (34), 26 variants of 50-residue mesophilic homologue BBL (35), 11 variants of 50-residue hYAP65 WW-domain (21, 36), and 10 variants of 50-residue villin HP35 (37, 38). The stabilities of these variants were measured previously. All sequences were assayed for sequence enrichment ratio defined as enrichment of a mutant before and after TMP-induced stability selection. As Figure 1b, 1c and Supplementary Figure S1 show that good correlations were observed for all proteins except that hYAP65 showed worse PCC values under stronger selection conditions. More specifically, at the condition of incubating *E. coli* with 25 μg/ml TMP for 2 × 12h (Figures 1b, Supplementary Figure S1a, S1c, S1e), Pearson Correlation Coefficient (PCC) values are 0.9, 0.58, 0.72, and 0.49 for Pin1 WW domain, BBL, villin and hYAP65, respectively. Strong correlations are robust under different selection pressures, which PCC values between 0.89 and 0.91 for Pin1 WW domain (Figure 1c). For BBL and villin HP35 longer incubation time and/or more concentrated TMP will lead to a slightly better PCC value (Supplementary Figure S1b, and S1d). However, PCC values for hYAP65 were low when treating the cells with 100 μg/ml or 400 μg/ml TMP, because a small library with a small number of mutants (10) are more susceptible to experimental noises with low counts of sequence numbers (Supplementary Table S1). Thus, the Sibs-Seq system preferentially selects more stable variants as intended.

### Initial test of Sibs-Seq using the uracil glycosylase inhibitor

To test if artificially generated structural homologs obtained from Sibs-Seq are useful for structural inference, we first selected a mixed helix/sheet protein: the uracil glycosylase inhibitor (a 83-residue protein of mixed helices and sheets, PDB ID: 6LYD) (39). To locate the best selection pressure for structure prediction, we performed experiments at 9 conditions with TMP concentrations ranging from 25 μg/ml, 100 μg/ml, to 400 μg/ml and incubation time ranging from 1 × 12 h, 2 × 12h, to 3 × 12 h. All variants selected from 9 conditions were sequenced. Each mutant’s stability is estimated from its fitness score according to the natural logarithm of its normalized sequence enrichment ratios (See Methods). As shown in Supplementary Figure S2, the PCC values between the fitness scores of the same mutants at different experimental conditions range from 0.72 to 0.94. Thus, the fitness score of a mutant as a stability indicator is robust across different experimental conditions.

As one example, Figure 2a displays the distributions of the fitness scores for single and double mutants, respectively, at the condition of incubating *E. coli* with 25 μg/ml TMP for 2 × 12h. (Only the distributions of single and double mutants are shown because there is only one mutant with >2 mutations as shown in Supplementary Table S2). The peak for the fitness distribution of single mutants at -0.13 is located only slightly below zero, the fitness score for the wild-type sequence, consistent with the expectation that the majority of single mutations have only minor effect on the protein stability. By comparison, the peak for the fitness distribution of double mutants is shifted to a lower fitness value to -0.48, also consistent with the expectation that changing more amino acid residues will likely be more destabilizing. Figures 2b, 2c show the distribution of the fitness scores along the sequence position. It indicates that low-fitness mutations are more like located at the interior rather than the surface of the protein, consistent with the fact that the mutations occurred at the structural core are more likely destabilizing (31). This result is further quantitatively supported by the correlation between the residue-wise average fitness score and residue solvent accessibility (RSA) with a PCC = 0.6 (Supplementary Figure S3). Figure 2c further shows the mutation coverage along the sequence position. The number of mutations at each position is ranging from 5 to 11, with the median number of 6. This indicates a high-quality mutant library generated by the error prone PCR in this work (See Methods).

**Figure 2.**
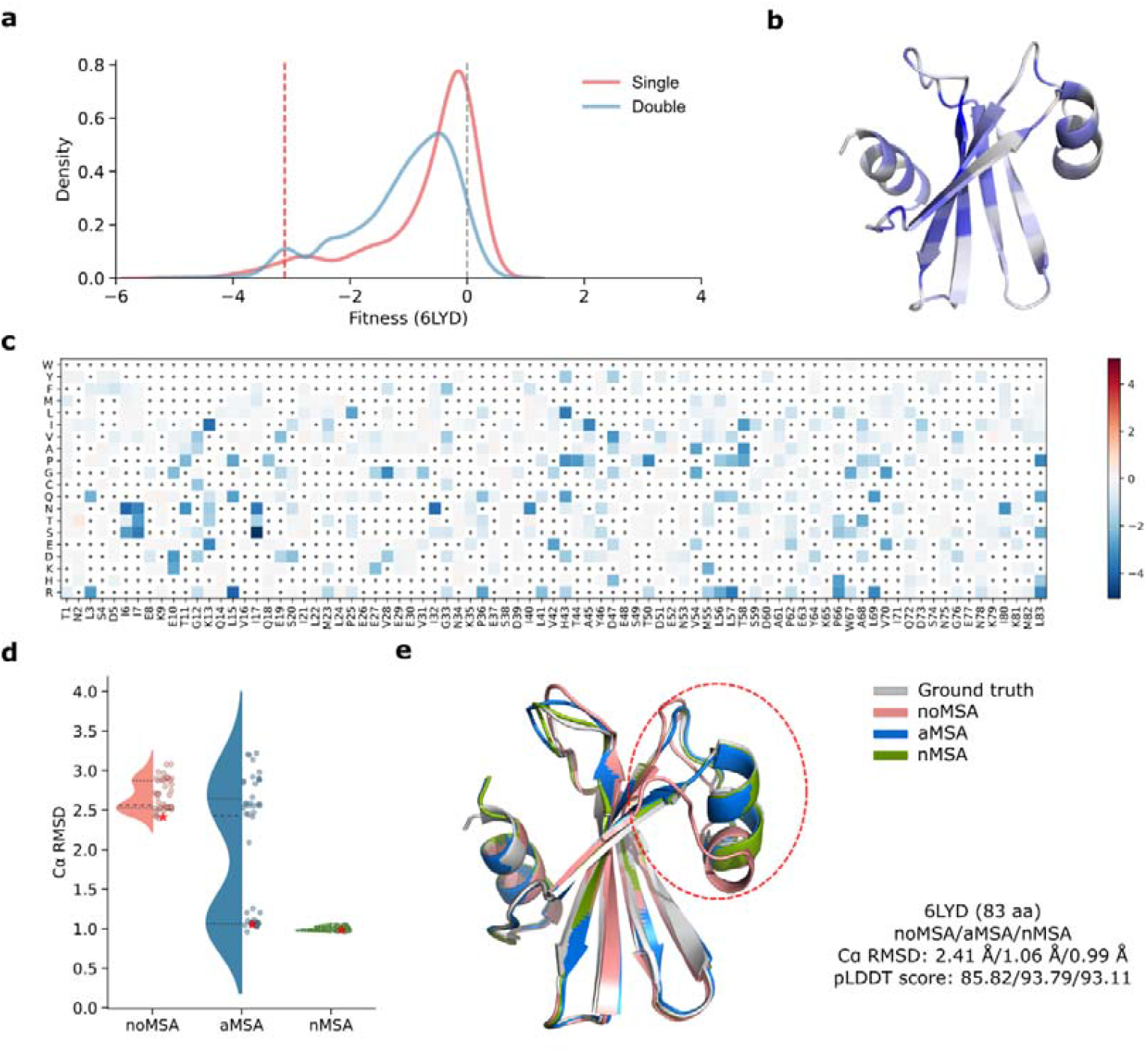
Enrichment of stable variants of the uracil glycosylase inhibitor with quality mutation coverage for high-accuracy structure prediction. **a**, Distributions of fitness scores for single and double mutants, consistent with the fact that double mutants are more destabilizing than single mutants (wild type has a fitness score at 0). Vertical dashed lines in grey and red indicate the fitness of the wild-type variants and the median fitness of Stop codon mutations, respectively. **b**, Domain structure colored by the average fitness values at each position, more negative fitness scores (darker blue) often associated with interior positions, consistent with the fact that the mutations at the structural core are more likely destabilizing. **c**, The heat map shows fitness values for single mutations observed on different sequence positions. White indicates wild-type stability, and red and blue indicate stabilizing and destabilizing mutations, respectively. Black dots indicate missing data. **d**, RMSD based on Cα atoms for predicted structures by AlphaFold2 without multiple sequence alignment (noMSA), with artificial MSA (aMSA) from Sibs-Seq, and with naturally occurring MSA (nMSA) as labeled. Star indicates the location of the structure predicted according the pLDDT score. **e**, The comparison between predicted structures without MSA (pink), with aMSA (blue) and nMSA (green) to the experimental structure (shown in gray, PDB ID 6LYD Chain B). Structures are predicted based on the highest pLDDT scores from 50 predictions. The largest region of the structural improvement from noMSA to aMSA or nMSA is circled. The data in this figure is based on the experimental condition of 25 μg/ml TMP for 2 × 12h’s selection.

### High-accuracy structure prediction for the uracil glycosylase inhibitor by AlphaFold2 with artificial homologous sequences

Predicting 3D structures by AlphaFold2 requires homologous sequences with the same stable structure. For convenience, we only employed mutants with a fitness score of no less than the wild type sequence (≥ 0). To obtain the variation of predicted structures, we ran 50 independent predictions. Figure 2d shows the RMSD values from native structures according to Cα atoms for 50 predictions without multiple sequence alignment (noMSA), with artificial homologous sequences generated by Sibs-Seq (aMSA), and with naturally occurring MSA (nMSA). Without MSA, AlphaFold2 provided a good prediction with RMSD between 2.39 Å and 3.08 Å but short of high-accuracy prediction (RMSD < 2 Å). Inputting aMSA will lead to a high-accuracy prediction for some runs. This high-accuracy prediction (1.06 Å) can be located by using the best pLDDT score, compared to 2.41 Å by the best pLDDT score, without MSA (Figure 2d). Compared with aMSA, the employment of nMSA led to a more consistent accuracy, with RMSD around 1 Å. There is essentially no difference between the predicted structure from aMSA and that from nMSA. The major improvement from noMSA to aMSA came from the correct prediction of the orientation of a short helix region (Figure 2e). Thus, for this protein, single and double mutations from aMSA are sufficient to make high-accuracy prediction (RMSD < 2 Å), whereas multiple mutations from nMSA can make a more consistent prediction.

Supplementary Figure S4 further illustrates the dependence of the accuracy on predicted structures at different selection pressures. In most cases, MSAs generated from either a longer TMP treatment or a higher TMP concentration gave a more consistent prediction accuracy, with median RMSD from 2.56 Å to around 1.1 Å (57% reduction) when using aMSAs from conditions (25-3, 100-1, 100-2, 400-1, 400-2, 400-3) for uracil glycosylase inhibitor. Supplementary Table S2 suggests that a longer incubation time leads to a smaller number of stable mutants, an indication of fewer false positives, whereas a higher TMP concentration leads to enrichment of more highly stable mutants (more true positives). Nevertheless, the final predicted structure according to pLDDT is near the same accuracy with RMSD ranging from 1.05 Å to 1.09 Å. The robustness of the result against different experimental conditions allows us to choose one condition for all subsequent experiments to save the time and the cost.

### High-accuracy structure prediction for the proteins of other structural topologies by Sibs-Seq and AlphaFold2

To confirm the generalizability of the above high-accuracy structure prediction, we randomly chose three proteins of different structural folds: a two-helix bundle (92-residue kinetoplastid membrane protein-11, KMP11, PDB ID: 7F0K), a four-helix bundle protein (97-residue human Arc C-lobe, PDB ID: 6TN7), and a mostly sheet protein (an 84-residue anti-CRISPR protein called AcrIE2, PDB ID: 7CHQ). For convenience, all proteins are referred by PDB ID, here and hereafter. PE250 HTS were employed for sequencing when the amino acid sequences of target proteins are greater than 88.

Supplementary Table S3 shows the statistics results of mutants before selection as well as after selection with ≥ 0 fitness scores at a given experimental condition. Substantially more single and double mutations are obtained than 6LYD due to the use of a larger mutant library. As for 6LYD (Figure 2a), the distribution of the fitness scores becomes broader, and shifted to lower fitness scores when the number of mutations changes from single, double to triple or more (Supplementary Figure S5), consistent with the expectation that changing more amino acid residues will more likely lead to less stable proteins.

Similar as 6LYD, the more exposed regions in these three tested proteins showed lower sensitivity to mutations (Supplementary Figures S6 and S7). It should be noted that most single mutations in 7CHQ do not have a detrimental impact on the folding stability of this protein. This might be related to the large flexible loop regions in this structural fold.

Figure 3a shows RMSD distributions of 50 predictions for 7F0K. For this simple structural fold of two-helix bundle, AlphaFold2 can already make a high-accuracy prediction with a narrow distribution and a 1.89 Å RMSD based on pLDDT without MSA. Adding aMSA generated from Sibs-Seq can even push the structural accuracy further down to 1.03 Å RMSD. This improvement is mostly due to a correct conformation of the N terminal end of the protein (Figure 3d). Interestingly, this is more accurate when compared with nMSA as nMSA failed to predict the structure with RMSD smaller than 2 Å. (Note: this protein was aligned in 86 residues due to missing 6 residues in its PDB structure).

**Figure 3.**
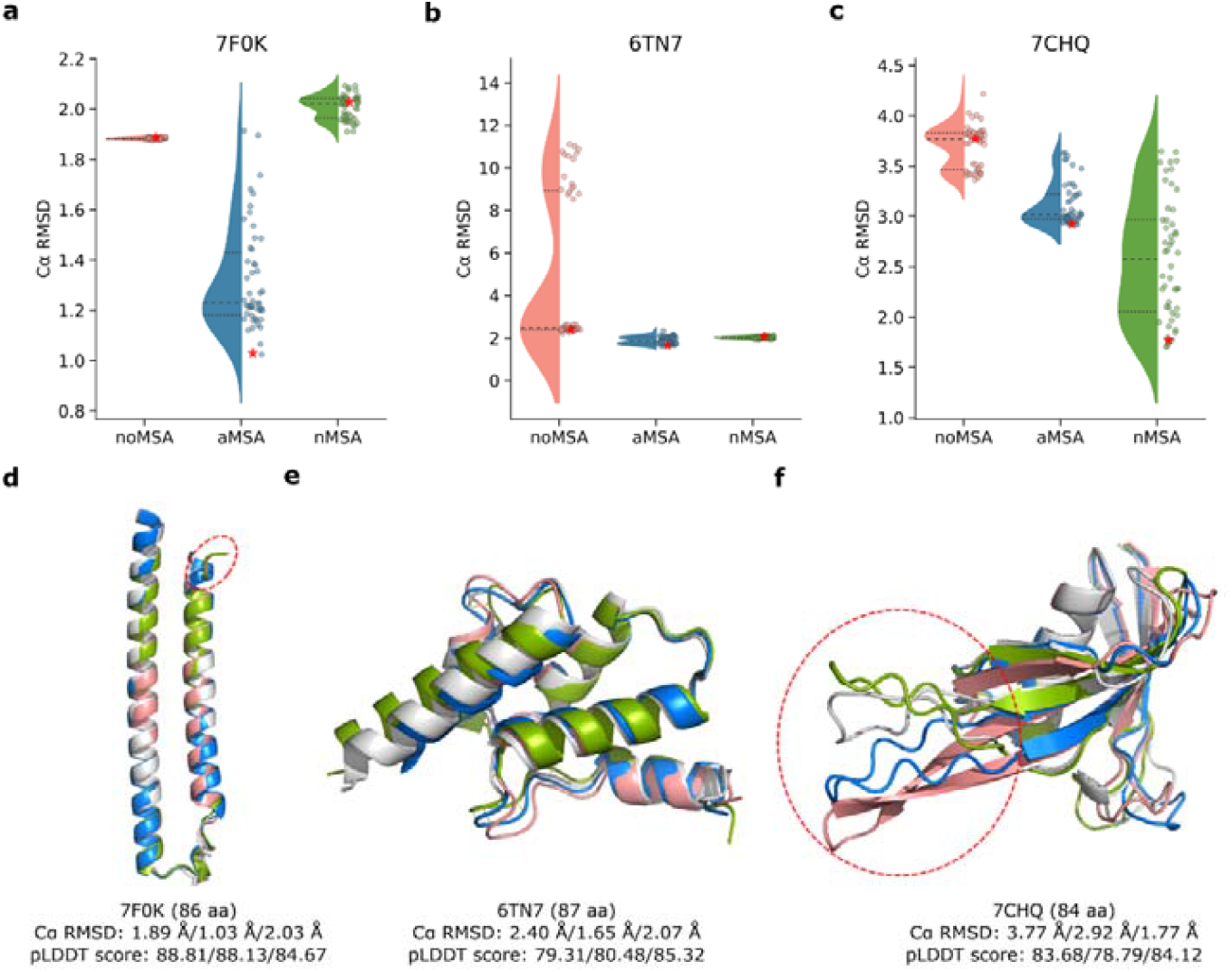
Accurate structure prediction by Sibs-Seq and AlphaFold2 for 7F0K, 6TN7, and 7CHQ. **a-c**, Comparison of 50 predicted structures in Cα RMSD at noMSA, aMSA and nMSA, respectively, with the prediction according to the highest pLDDT score shown in red star for 7F0K (a), 6TN7 (b) and 7CHQ (c). **d-f**, Predicted structures with the highest pLDDT scores from 50 predictions were aligned to the experimental structures for 7F0K (d), 6TN7 (e) and 7CHQ (f). Ground truth is shown in gray. The AlphaFold2 predictions with no MSA, aMSA, and nMSA are displayed in pink, blue and green, respectively.

Figure 3b shows the result for 6TN7. For this four-helix bundle, AlphaFold2 can make a good prediction with a 2.40 Å RMSD based on pLDDT without MSA. Adding aMSA generated from Sibs-Seq can even push the structural accuracy further down to 1.65 Å RMSD which is slightly lower than nMSA (2.07 Å). It should be noted that noMSA failed to get the correct fold for some runs. The utilization of aMSA and nMSA all led to a more consistent accuracy. (Note: this protein was aligned in 87 residues due to missing 10 residues in its PDB structure).

Figure 3c shows the result for 7CHQ. For this mostly sheet protein, AlphaFold2 can make a reasonable prediction with a 3.77 Å RMSD based on pLDDT without MSA. Adding aMSA generated from Sibs-Seq can even push the structural accuracy further down to 2.92 Å RMSD. This improvement is largely due to the improvement of the extruded β-hairpin region as shown in Figure 3f. Despite of this extruded β-hairpin region, using nMSA was able to further improve the prediction accuracy to 1.77 Å. This special case (the only one with > 2 Å RMSD) will be discussed more later.

### High-accuracy structure prediction for an ‘orphan’ protein SeviL

We further apply Sibs-Seq to an ‘orphan’ protein called SeviL (PDB ID: 6LF2) studied previously (9). SeviL is a GM1b/asialo-GM1 binding R-type lectin which adopts a β-trefoil fold consisting of three sub-domains that each contains four β-strands. For this protein, we obtained 10586 mutants before and 1317 mutants after the selections (Supplementary Table S4). Supplementary Figure S8 shows that the fitness scores of different mutants of the orphan protein SeviL. Unlike the single mutants of the other 4 tested proteins, most single mutants of SeviL showed fitness lower than the wild type sequence, indicating that this structure is more sensitive to mutation. The correct trend of less stable variants for more mutations is observed. The higher sensitivity to mutations should be related to the more complex structure of SeviL. The distribution of fitness scores along the sequence position (Figure 4a and 4b) also shows that most of the structure is sensitive to mutations except for few sites at the terminal or surface. Correlation between the fitness scores and solvent accessibilities could also be observed in this protein, with PCC value of 0.38 (Figure 4c).

**Figure 4.**
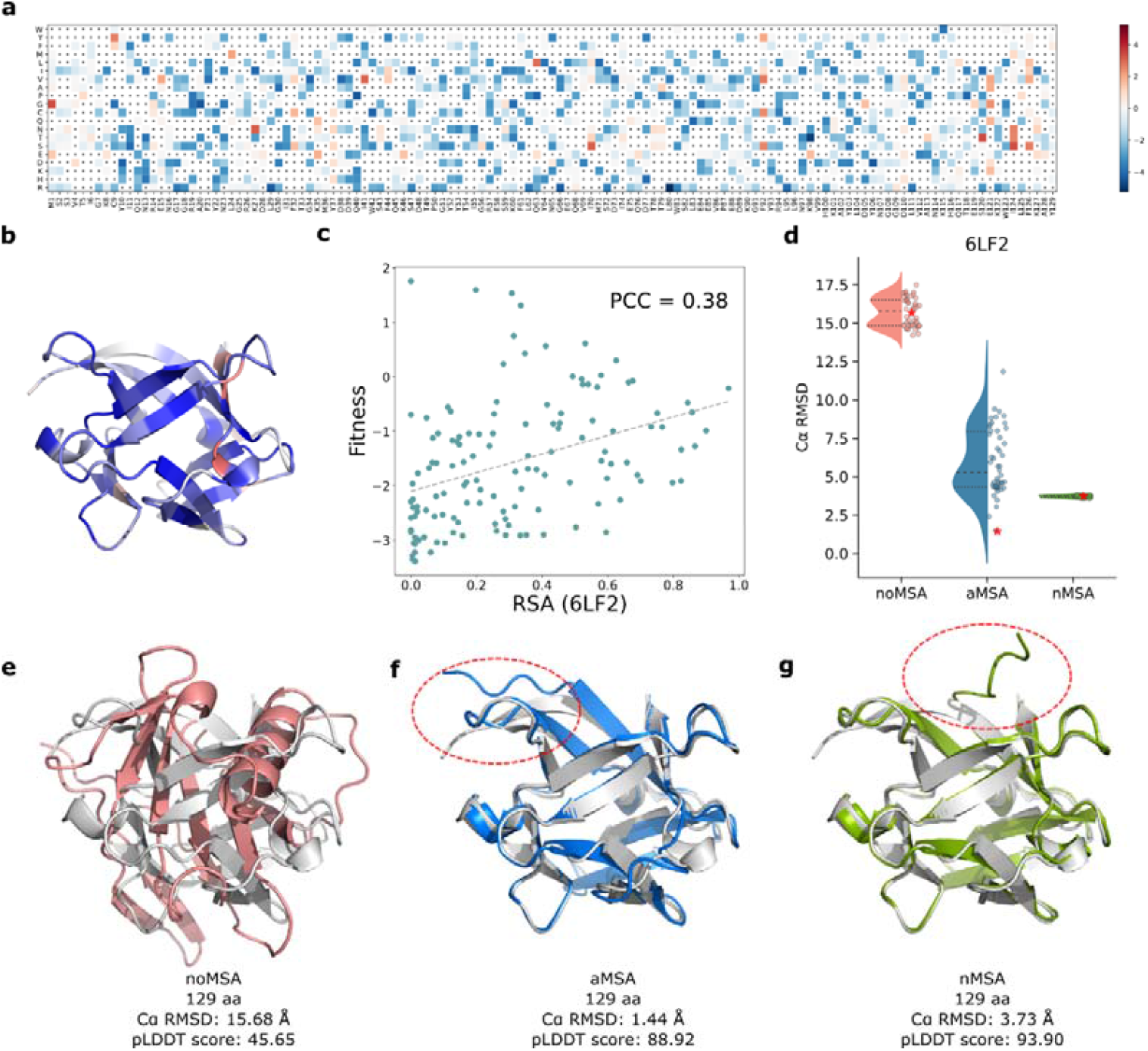
High-accuracy structure prediction of a 129-residue ‘orphan’ protein SeviL (PDB ID: 6LF2). **a**, The distribution of fitness values on different positions of SeviL. The heat map shows fitness values for single mutations observed on different positions. White indicates wild-type stability, and red and blue indicate stabilizing and destabilizing mutations, respectively. Black dots indicate missing data. **b**, Distribution of fitness scores along the positions of SeviL. **c**, Correlation between the residue-wise average fitness score and the relative solvent accessibility (RSA) for 6LF2. **d**, Comparison of the predicted structures in Cα RMSD given by AlphaFold2 with noMSA, aMSA, and nMSA. **e-g**, Predicted structures with the highest pLDDT scores from 50 predictions were aligned to the experimental structure. Ground truth is shown in gray. The AlphaFold2 predictions with noMSA, aMSA, and nMSA are displayed in red, blue, and green, respectively. The most significant improvement of the use of aMSA over the use of nMSA is circled in red.

Figure 4d compares the accuracy for the structures predicted by AlphaFold2 with noMSA, aMSA and nMSA. AlphaFold2 is unable to predict the structure of this protein without MSA (RMSD > 16 Å). The incorporation of artificial MSA is sufficient to yield the best predicted structure at high accuracy of 1.44 Å RMSD (according to pLDDT). This performance was even better than nMSA, which achieved prediction accuracy at 3.73 Å RMSD. The major improvement came from the correct prediction of the N terminal region of the protein (Figure 4e, 4f).

### Improved structure prediction for a CASP16 target T2284

In addition, we employed the same strategy to study a protein target used in the 16^th^ CASP (Critical Assessment of Techniques for Protein Structure Prediction). The protein target T2284 (gp62 of P23-45 phage) contains 120 amino acids in total and has no published structural information. We obtained 4067 mutants before and 1413 mutants after the selections for this protein. Unlike the other 5 tested proteins, T2284 shows lower overall sensitivity to mutations, consistent with smaller differences in fitness scores of different mutants (Figure 5a and 5b). What is more interesting is that mutations at several positions show significantly improved stability in T2284 (Figure 5a).

**Figure 5.**
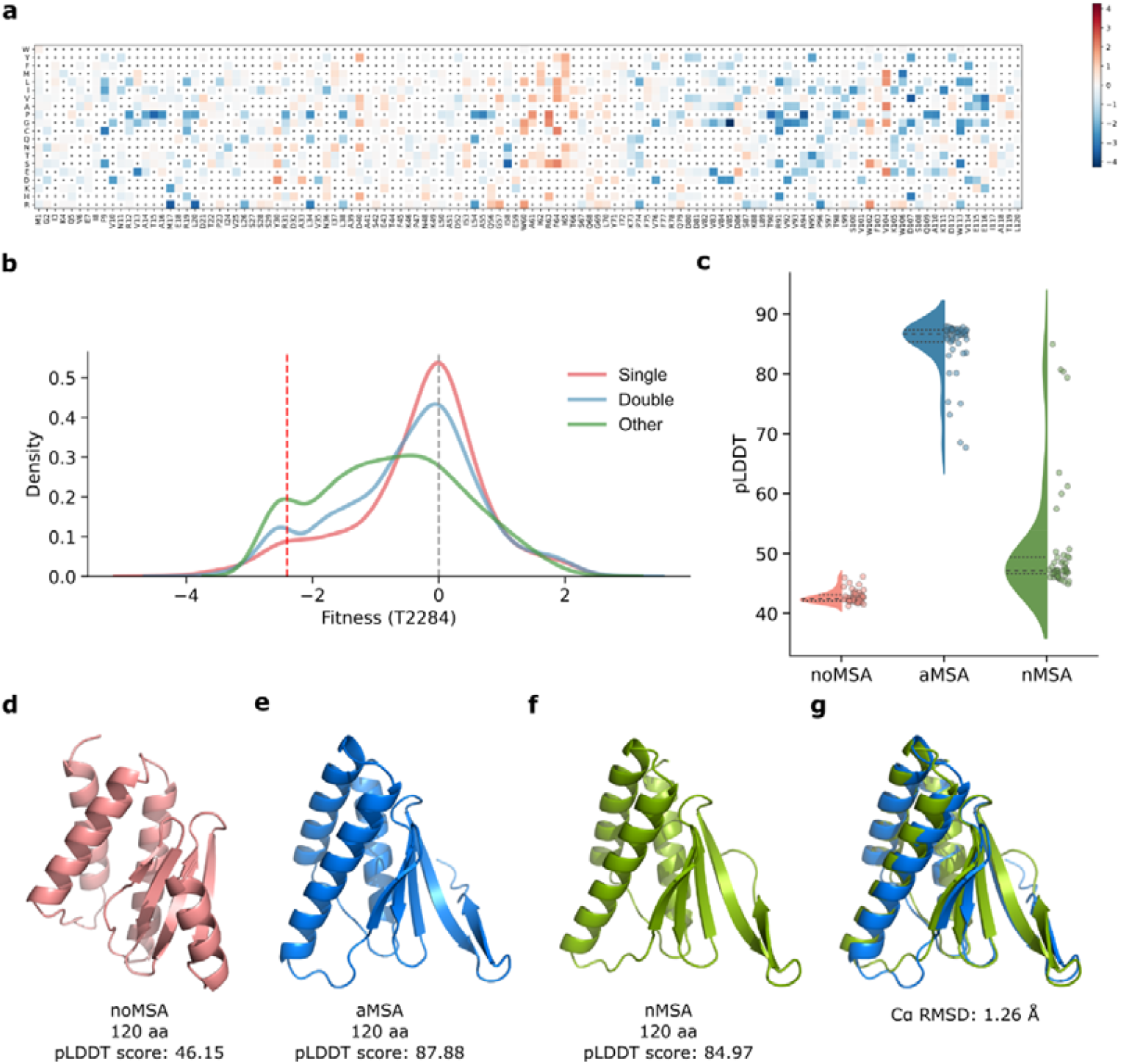
High-accuracy structure prediction of a 120-residue CASP16 target T2284. **a**, The distribution of fitness values on different positions of T2284. The heat map shows fitness values for single mutations observed on different positions. White indicates wild-type stability, and red and blue indicate stabilizing and destabilizing mutations, respectively. Black dots indicate missing data. **b**, Distribution of the fitness scores for single, double, and other mutants for T2284 is consistent with larger damaging effect on protein stability with more mutations. Vertical dashed lines in grey and red indicate the fitness of the wild-type variants and the median fitness of Stop codon mutations, respectively. **c**, Comparison of the predicted structures in pLDDT score given by AlphaFold2 with noMSA, aMSA, and nMSA. **d-f**, Predicted structures with the highest pLDDT scores from 50 predictions. The AlphaFold2 predictions with noMSA, aMSA, and nMSA are displayed in red, blue, and green, respectively. **g**, Comparison of top predicted structures with aMSA and nMSA.

Figure 5c compares the accuracy for the structures predicted by AlphaFold2 with noMSA, aMSA and nMSA. Without using MSA, AlphaFold2 is unable to predict the structure of this protein with high confidence (pLDDT < 50). The incorporation of artificial MSA significantly improved the confidence of predicted structures, even better than the performance of nMSA (Figure 5c-f). The best predicted structures of aMSA and nMSA achieved a pLDDT score of 87.88 and 84.97, respectively. The major difference between these two structures came from the flexible loop regions of the protein, with a RMSD of 1.26 Å (Figure 5g). These two structures (Figure 5e-g) all indicate that T2284 adopts a structural fold consisting of a 3-helix bundle and one β-sheet.

## Discussion

The success of AlphaFold2 for mapping natural homologous sequences to a single structure in high accuracy raises the question how to predict the structures for those proteins with insufficient number or coverage of naturally occurring mutations. Here, we proposed to generate artificial homologous sequences by developing a structure-stability-based mutational scanning assay, followed by high-throughput sequencing (Sibs-Seq). The stability-based selection was validated by the high correlation observed between measured unfolding stabilities and sequence enrichment ratios of mutants of four proteins (Figure 1 and Supplementary Figure S1). Artificial homologous sequences generated by Sibs-Seq allow AlphaFold2 to make high-accuracy structure prediction (RMSD < 2Å) for four of the five proteins tested. More importantly, a single round of mutations generated by error-prone PCRs (i.e., without directed evolution) is sufficient to make such a high-accuracy prediction.

The only exception is a protein with fully exposed extruded beta-hairpin (7CHQ), which is constrained in a crystal packing environment but unlikely constrained as a monomer in a solution (Supplementary Figure S9).

We developed the mDHFR-based tripartite fusion system for structure-stability-based selection. Other biosensors were also available to detect protein stability *in vivo* (40, 41). One conceptually similar, commonly-used system is a β-lactamase-based biosensor (25). However, indirect resistance of β-lactamase (42) may lead to a large number of false-positive clones, interfering with the screening results of β-lactamase biosensors. Moreover, unlike cytoplasmic mDHFR, the β-lactamase is located at the outer membrane-bounded periplasmic space, which makes it unsuitable for screening the stabilities of proteins containing multiple reduced cysteines. Another stability-based selection employed in *E. coli* is the CysG^A^ tripartite system (28), which utilizes the intensity of red fluorescence to measure the stabilities of proteins *in vivo*. The current form of the technology is not yet amenable for high-throughput screening. Moreover, the CysG^A^ biosensor needs weakly stable variants as the template to start (28), compared to the direct use of wild-type in the mDHFR system developed here. Other library□based display technologies were also utilized for protein stability studies (43). For example, the recent two studies employed protease cleavage in combination with yeast or cDNA display to analyze the folding stabilities of natural and de novo designed proteins (23, 24). These two methods, however, are not suitable for proteins of > 72 amino acids due to limitation associated with proteolysis (23, 24). By comparison, the mDHFR biosensor developed here allows an easy integration with high-throughput sequencing.

The enriched structure-stable, artificial homologous sequences with minimal mutations allow us to demonstrate their usefulness for structural inference. In addition to its utility for collecting structure-stable monomeric variants in this work, the mDHFR tripartite biosensor can be easily extended to protein-protein binding (30, 31) and optimization of de novo designed proteins through directed evolution (44). The work in this area is in progress.

One surprising result from this study is that one single round of random mutations, selected by the mDHFR stability sensor and enriched for 24 or 36 h incubation, is sufficient to allow AlphaFold2 to make high-accuracy prediction (RMSD < 2 Å), even for the protein 6LF2 that AlphaFold2 is unable to find its correct fold with the single sequence input. For the case of 83-residue 6LYD, only 79 single and 100 double mutations are all what is needed for high-accuracy prediction (Supplementary Table S2). For the challenging orphan protein (6LF2), 1317 unique sequences are sufficient to yield high-accuracy structure prediction (Supplementary Table S4). Thus, the technique developed here should be applicable in principle to the proteins larger than the 129-residue 6LF2 tested here. A limit in length (600 nt, 200 aa for proteins) by second-generation sequencing techniques can be resolved by the use of the third-generation sequencing platforms (45). For example, the ampicillin-selection of ampicillin-resistant β-lactamases genes allowed deep mutational scanning and sequencing of > 260 residues PSE1 and TEM-1 β-lactamases by the third generation sequencing platform (Pacbio) (17, 18).

It should be noted, however, that naturally occurring homologous sequences, with more evolution information from accumulation of multiple mutations at the same time, provide more consistent prediction (narrow distribution in RMSD) in 4 out of 5 proteins studied here. This consistent prediction from nMSA with low sequence identities, however, could also be because AlphaFold2 was trained by nMSA. Re-training AlphaFold2 with aMSA with high sequence identity (when sufficient data is available) may address this problem.

Nevertheless, integration of Sibs-Seq with AlphaFold2 provides an inexpensive, and efficient alternative to NMR, X-ray crystallography or cryo-EM for those proteins that AlphaFold2 failed to provide a high-accuracy prediction. One notable advantage over structure determination by NMR, X-ray crystallography or cryo-EM is that Sibs-Seq can be conducted in a molecular biology laboratory without the need for any expensive equipment. Moreover, for a typical protein of <200 residues, the consumables will cost less than 1000 US dollars, including gene synthesis, EP-PCR, T4-DNA ligase, PE250 HTS sequencing and etc (< 300 US dollars for PE150 HTS sequencing of proteins < 90 residues). The overall time required for library construction, TMP-based selection, and HTS sample preparation for processing several proteins simultaneously is < 3 weeks for a person skilled in routine molecular biology experiments. Additional times for gene synthesis, outsourcing HTS and data processing will lead to a time frame of solving the structures of multiple proteins simultaneously within two months. Streamlining the workflow, automating some wet-lab experiments and in-house high-throughput sequencing can further cut the time requirement down. For example, one can purchase a synthesized doped library, rather than EP-PCR. Currently, two or more rounds of EP-PCR may be performed to control the mutation rate, according to Sanger sequencing results. This would take several working days. On the other hand, a synthesized doped library can be preset with a given mutation rate, and thus, speed up the whole process.

Finally, we noted that the Sibs-Seq technique in the current form is limited to soluble and nontoxic proteins foldable in *E. coli*. These issues may be addressed by transferring the mDHFR-based stability assay to other host cells in which the target protein is foldable.

## Materials and Methods

### Design of constructs

All DNA sequences used in this research are listed in Supplementary Table S5. The split murine DHFR gene (SmD) was designed with one AmpR promotor at its 5’ site and two (GGGGS)_2_ linkers with KpnI and BamHI restriction endonuclease sites between the two SmD gene fragments mF1,2 and mF3. The SmD sequence (29, 33) was directly synthesized (General Biosystems, China) and cloned between the PciI and NdeI restriction sites of the backbone vector pUC57-Kan (GenScript).

### Selection of natural proteins for mutational scanning

We selected five natural protein structures with different folds from PDB that were released after May 2020 (the start date of CASP14) and a CASP16 target T2284. Structures with <50 or with simple topologies (for example, a single α-helix) were removed. Proteins with more than 150 amino acids were not considered in this study, so that we can focus on the use of the low-cost second-generation sequencing technique.

### Construction of the mutant library

The 108-oligo pool with KpnI and BamHI restriction sites was synthesized by Integrated DNA Technologies. It was amplified using PrimerSTAR HS DNA polymerase (Takara), digested with restriction enzymes, and then cloned into the pUC57-SmD vector by using T4 DNA ligase (Takara). The pUC57-SmD-108 constructs were employed to evaluate the correlation of protein folding stability with TMP resistance in our selection system.

For five selected protein targets, we employed error-prone PCR (EP-PCR) to generate the mutated DNA libraries (46). The random mutagenesis kit was purchased from Beyotime, with Evo-FP and Evo-RP as the primers. EP-PCR products were then separated by using 5% Urea-PAGE, extracted, and finally purified using ethanol precipitation, similar to our previous study (16). The purified products were cloned into the pUC57-SmD vector by using T4 DNA ligase. The ligated products were purified and eluted with nuclease-free water for electro-transformation. Two or more rounds EP-PCR were performed, the recycled target DNA products were cloned into pUC57 vector and transformed to DH5α for library construction. The colonies were randomly picked for Sanger sequencing to calculate the mutation rate.

Preparation of the electrocompetent DH5α cells and electroporation were done following the instruction manual of the micropulser electroporator (Bio-rad). After electroporation, the *E. coli* transformants were then cultured in 20 ml LB medium for 1 h, with the addition of Kanamycin to the final concentration of 50 μg/ml for additional cell growth (37 °C, 10 h). The overnight cell culture was pooled and resuspended in LB medium containing 15% glycerol and stored at -80 °C. The sizes of constructed libraries were estimated by counting the number of colonies after gradient dilutions of the fresh transformants on LB agar plates with kanamycin.

### TMP selection

20 μl frozen stocks from the diluted library (10^6^10^7^ clones) were transferred to 20 mL EZ Rich Defined Medium containing 0.1% glucose and 50 μg/ml kanamycin. The cells were cultured at 37 °C overnight with shaking at 200 rpm. For the first round of selection, 20 μl overnight cell culture was transferred to 20 ml EZ Rich Defined Medium containing 0.1% glucose, 50 μg/ml kanamycin, and a specific concentration of TMP for additional cell culture (12 hours at 37 °C). Meanwhile, the remaining cell culture was harvested by centrifugation and the pellets were stored at -20 °C for plasmid extraction.

### Sample preparation for next-generation sequencing

All the cell pellets were thawed and the plasmids were extracted by using Plasmid Mini Kit I (Omega Bio-tek). The plasmids were then employed as the templates for sequencing library constructions. The PCR products with sequencing adapters were gel-purified and sent to Novogene for next-generation sequencing. All the libraries were sequenced on NovaSeq 6000 in the paired-end mode.

### Processing of next-generation sequencing data

Paired-end fastq files from next-generation sequencing were merged by using PEAR (47). Both 5’ constant regions and 3’ constant regions of the assembled reads were trimmed by cutadapt (48). Then we used FASTX-Toolkit (http://hannonlab.cshl.edu/fastx_toolkit/) and vsearch (49) to filter sequencing reads with low quality, and count the read number of identical sequences. The read numbers before and after selection were then summed up for each unique sequence. The enrichment ratio of a DNA variant was calculated as the ratio of output (after selection) to input (before selection) read count, i.e.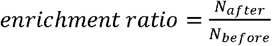, with *N* as read counts, subscripts denoting the sequencing library before or after TMP selection. The read counts were further processed by the STEAM module from DiMSum (50) with minor adjustments. DiMSum STEAM performed statistical analyses to obtain the fitness score and error estimates for each protein variant. The fitness score of a protein variant was calculated as the natural logarithm of normalized enrichment ratio, i.e. 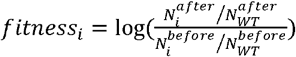, with superscripts denoting the sequencing library before or after TMP selection, subscripts denoting protein variant i or the wild-type variant. In this definition, the fitness score of the wild-type protein is 0. A protein variant with a positive fitness score, is likely to have a higher folding stability than the wild-type protein, whereas a negative fitness score indicates a variant likely with weaker stability than the wild type.

### The overall strategy for generating artificial MSAs from sequencing data

Stable homologs were selected according to the fitness scores and error estimates. This is accomplished by in-house Python scripts. The cutoff for fitness score was set to 0 by default.

### Structural modeling by AlphaFold2

AlphaFold2 (v 2.2.0) was installed and separated into two individual steps. The first step was the preprocessing of the input sequences, with the pickle files containing features as the output. All the remaining processes made the second step. In order to incorporate artificial homologous sequences, we skipped the first step and used household Python scripts to generate the pickle file as the input for the second step. AlphaFold2 were run for three cases: noMSA (no MSA employed), aMSA (artificial MSA from Sibs-Seq), and nMSA (natural MSA). Natural MSA was obtained from the homology search result of AlphaFold2 (v2.2.0). For all cases, no structural homologs were included for this study to avoid the use of homologous templates for prediction. We ran 50 independent predictions for each case and every protein. We selected the top model based on the predicted local distance difference test score (pLDDT) for each prediction.

### Statistics and reproducibility

For the experimental part, we did not perform multiple experiments under the same conditions. But for the 108-oligo pool and one natural protein (PDB ID: 6LYD), we used different TMP concentrations and different incubation times for selection of stable variants to confirm reproducibility. For structure prediction, we ran prediction independently for 50 times and all 50 models were used for downstream analysis.

## Data availability

Folding stability measurements of the 108-oligo pool can be directly downloaded from the Supplementary materials of a previous study (23). All original sequencing data have been deposited in the Gene Expression Omnibus with accession number GSE248664. All processed data are available for download at https://doi.org/10.5281/zenodo.10199203.

## Acknowledgement

This work was supported by the National Key Research and Development Program of China (no. 2021YFF1200400) and the supercomputing facility of Shenzhen Bay Laboratory.

## Author Contributions

JZ and JT conducted experimental design. JT performed wet-lab experiments. ZZ performed data analysis, computational prediction, and method development. JZ and YZ initiated and supervised the project. YZ provided the funding support. YZ, ZZ, and JT drafted the initial manuscript. All authors involved subsequent manuscript improvement.

