## Supplementary materials for "Effective High-Accuracy Prediction of Protein Structures from Easily Obtainable Artificial Homologous Sequences by Structure-Stability-Based Selection"

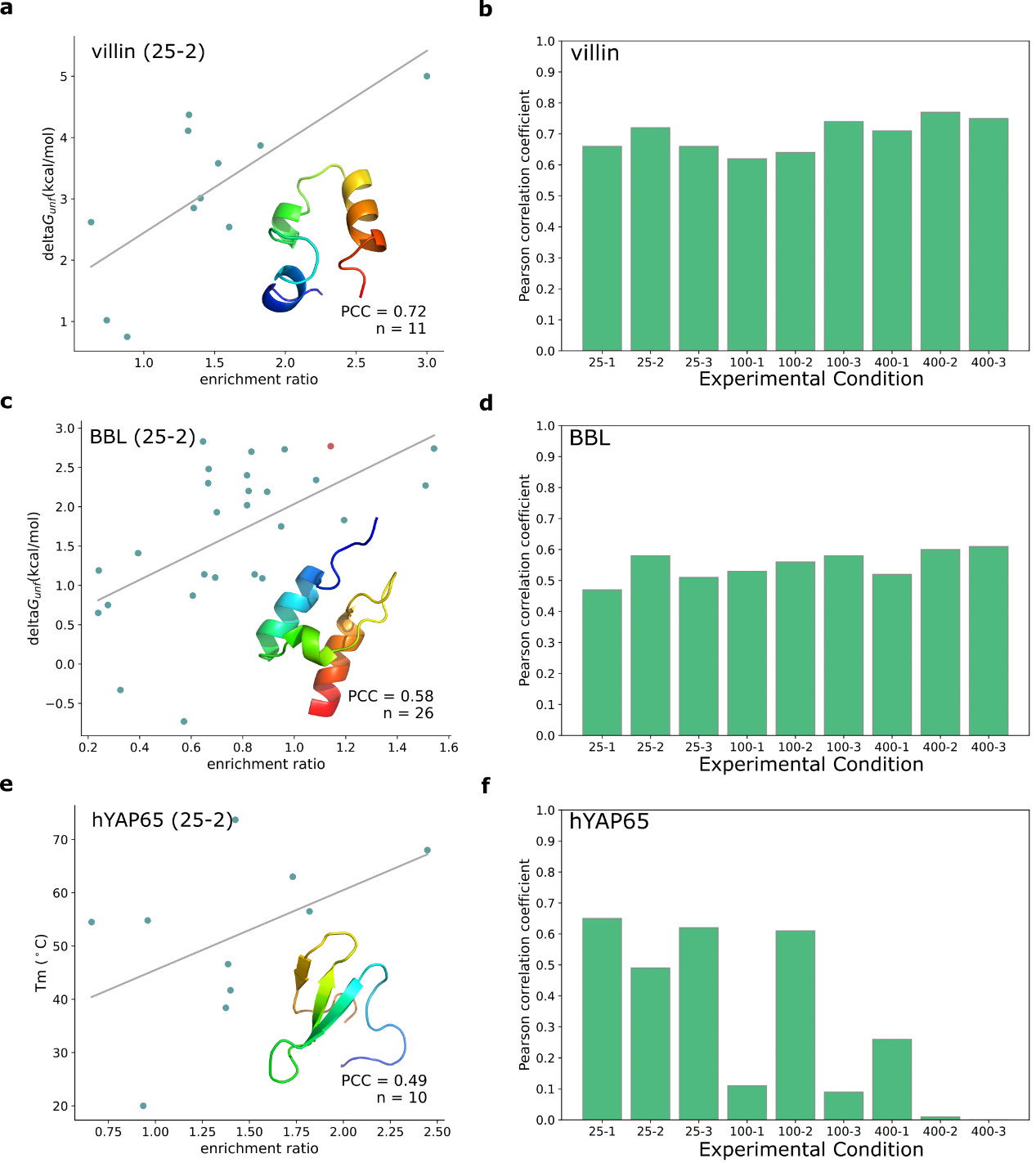
 **Supplementary Figure S1. Free energy of unfolding is correlated with enrichment of mutation variants in Sibs-seq.** In addition to Pin WW domain, the Sibs-Seq assay was also validated by using 11 variants of villin (**a, b**), 26 variants of BBL (**c, d**), and 10 variants of hYAP65 (**e, f**) with known stabilities (ΔG_un_ in kcal/mol or Tm in ℃). We used BBL ΔG_unf_ data measured by thermal denaturation, villin HP35 ΔG_unf_ data measured by urea denaturation, and Tm data for hYAP35. **a**, **c**, and **e** show the relation between ΔG_unf_ of a mutant and its sequence enrichment ratio at the experimental condition of 25 μg/ml TMP for 2 × 12h with Pearson Correlation Coefficient, (PCC)=0.72, 0.58, and 0.49 for villin, BBL, and hYAP65, respectively. The structures were drawn according to PDB ID: 1VII for villin, 1W4H for BBL, and 1K9Q for hYAP65. The red dot represents the position of the wild-type sequence if available. **b, d,** and **f** display PCC values observed for all selection pressures tested (TMP concentrations from 25 μg/ml, 100 μg/ml, to 400 μg/ml and incubation time from 1 × 12 h, 2 × 12 h, to 3 × 12 h) for villin, BBL, and hYAP65, respectively.

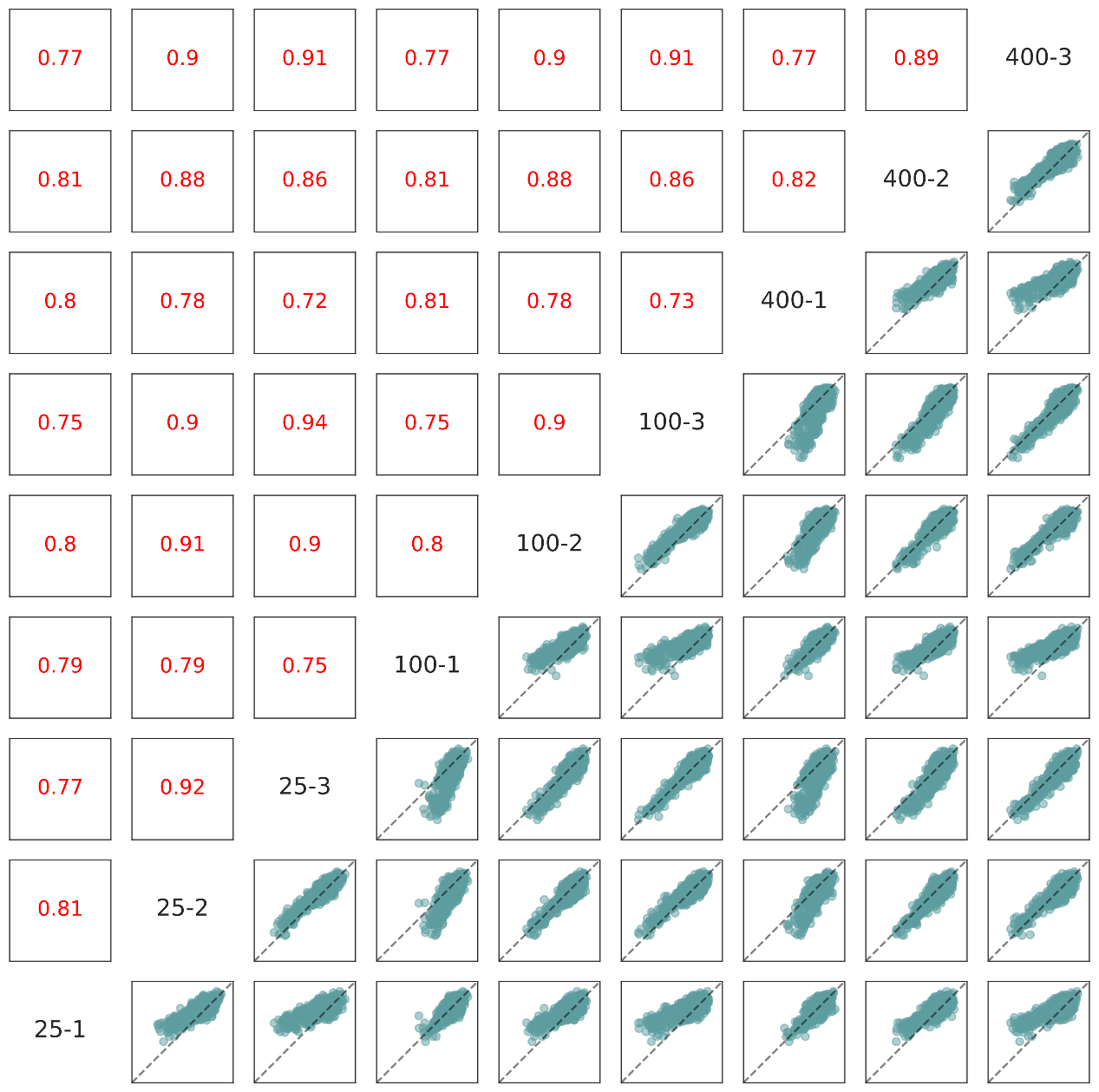

**Supplementary Figure S2. Consistency between fitness scores of the same mutant at different experimental conditions.** Pearson correlation coefficients (upper triangles) and scatter plots of the fitness scores of the same mutant at different experimental conditions as labelled. The first number indicates the concentration of TMP in μg/ml and the second number indicates the duration of TMP incubation in number of 12 h.

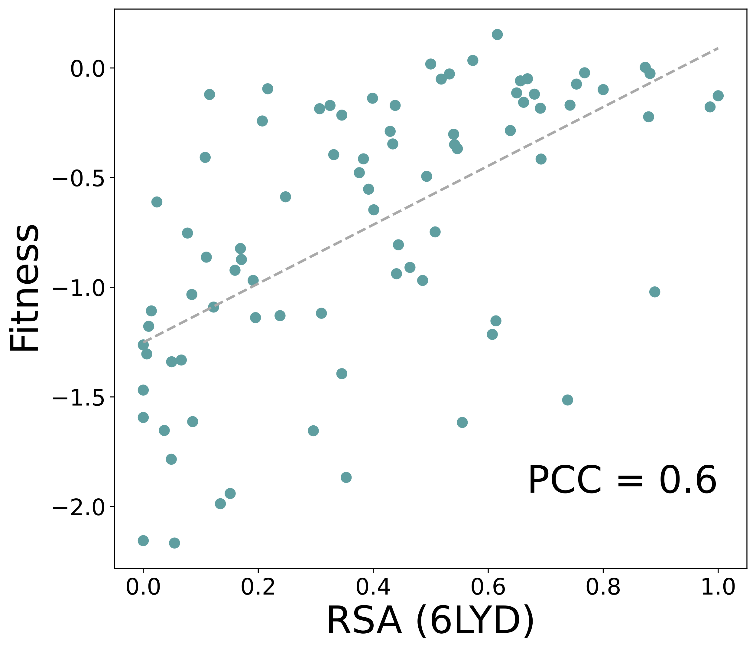

**Supplementary Figure S3. The average fitness score at each sequence position is correlated with its relative solvent accessibility (RSA) for 6LYD.** Pearson Correlation Coefficient (PCC) is 0.6 as shown.

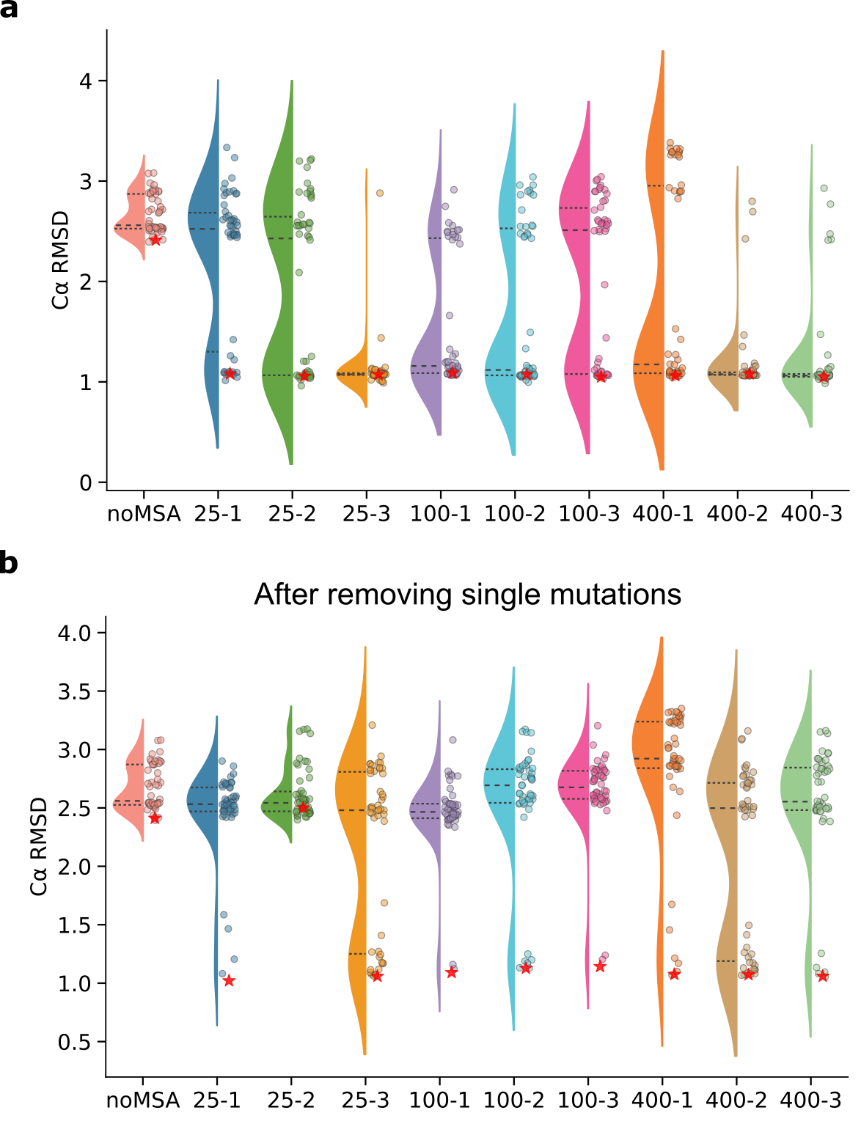

**Supplementary Figure S4. Comparison of structure models (6LYD) predicted with MSAs from different experimental conditions.** noMSA indicates the prediction made with single sequence as input only. The first number in the X-axis label indicates the concentration of TMP in μg/ml and the second number indicates the duration of TMP incubation in number of 12 h.

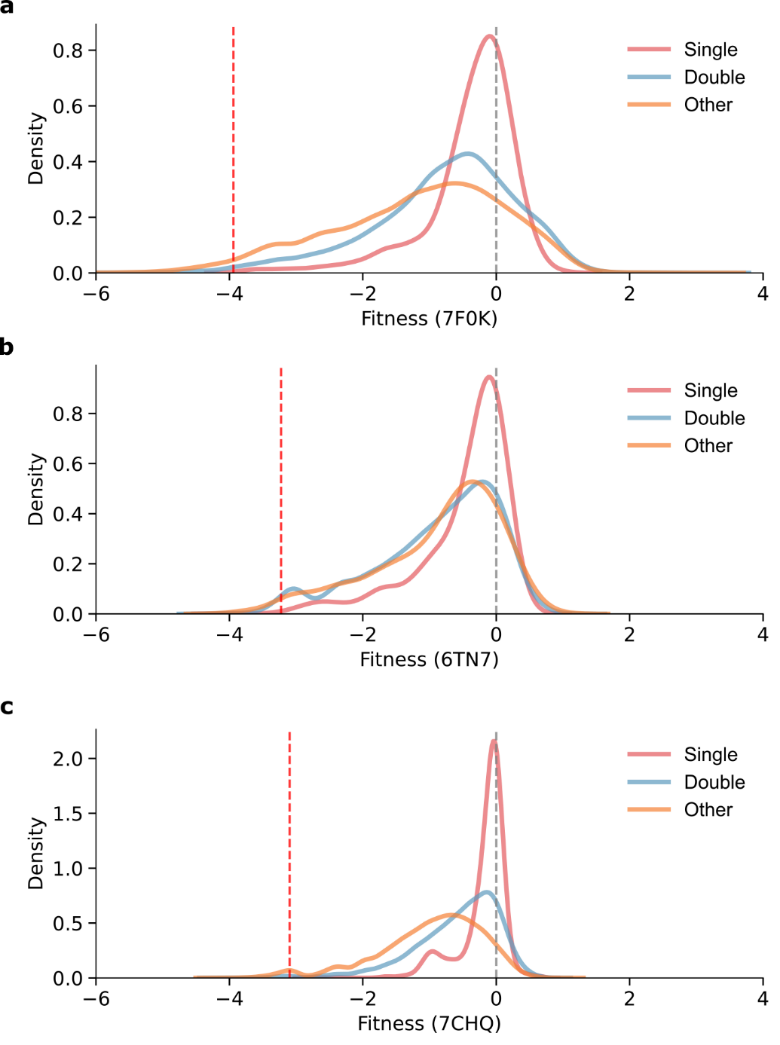

**Supplementary Figure S5. Distributions of the fitness scores for single, double, and other mutants for 7F0K, 6TN7, and 7CHQ** **from top to bottom are consistent with larger damaging effect on protein stability with more mutations**. Vertical dashed lines in grey and red indicate the fitness of the wild-type variants and the median fitness of Stop codon mutations, respectively.

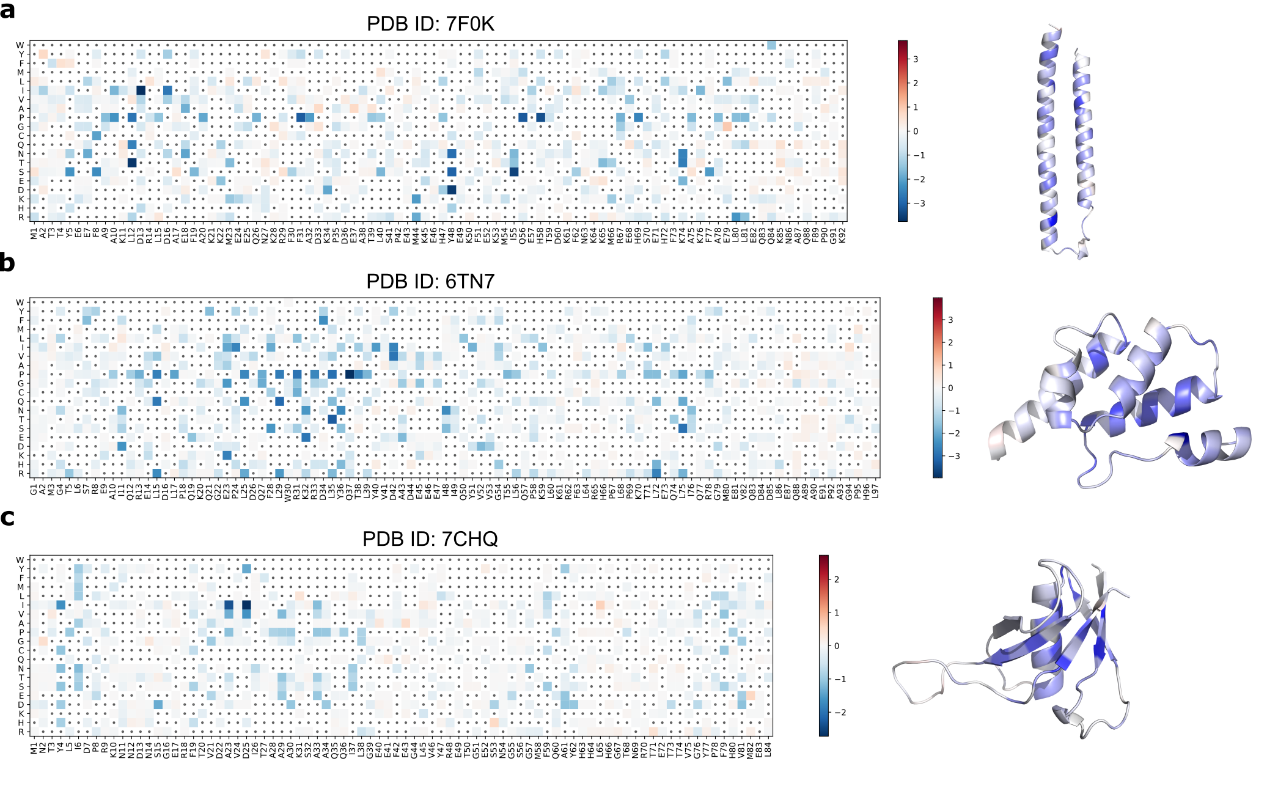

**Supplementary Figure S6. The distributions of fitness values on different positions of three target proteins: 7F0K (a), 6TN7 (b), and 7CHQ (c) indicate unbiased quality mutation libraries.** Left, the heat map shows fitness values for single mutations observed on different positions. White indicates wild-type stability, and red and blue indicate stabilizing and destabilizing mutations, respectively. Black dots indicate missing data. Right, domain structure colored by the average fitness values at each position. Darker blue indicates that mutations are more destabilizing.

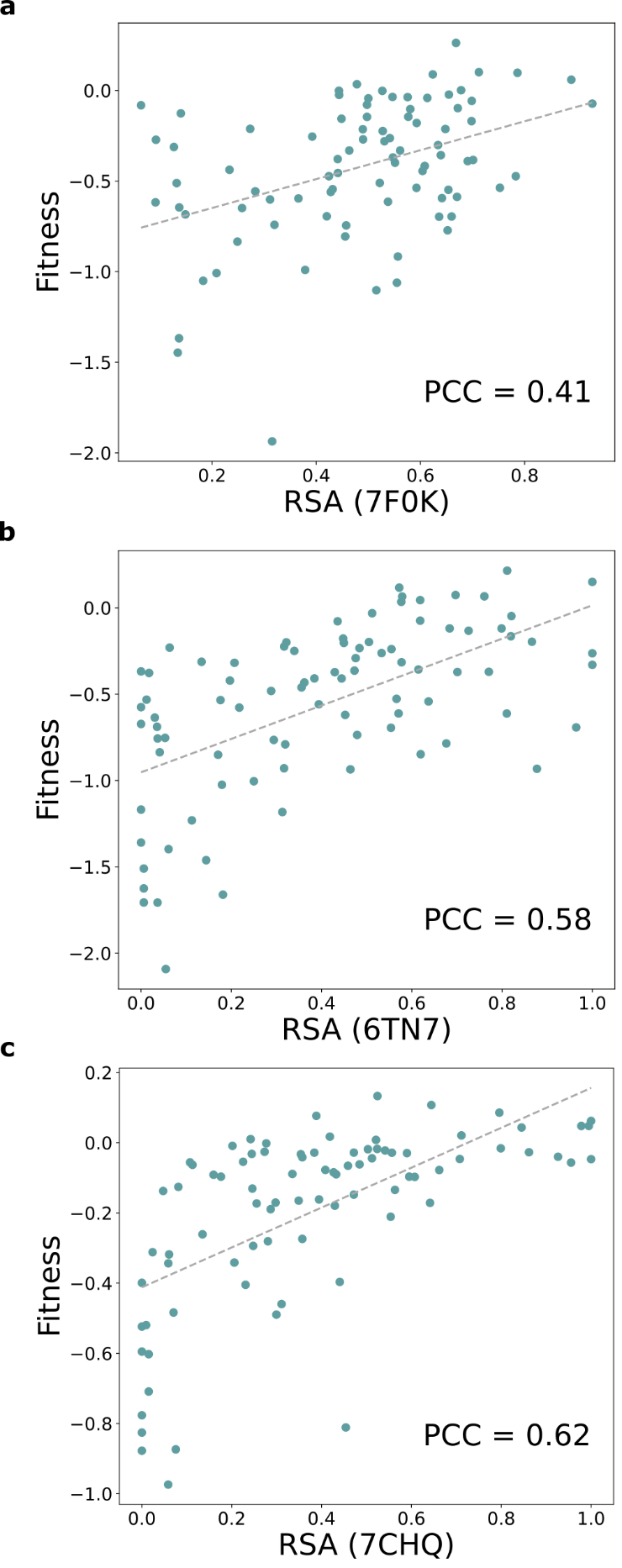

**Supplementary Figure S7. The average fitness score at each sequence position is correlated with its relative solvent accessibility (RSA) for 7F0K (a), 6TN7 (b), and 7CHQ (c).** Pearson Correlation Coefficient (PCC) values are indicated in the figure.

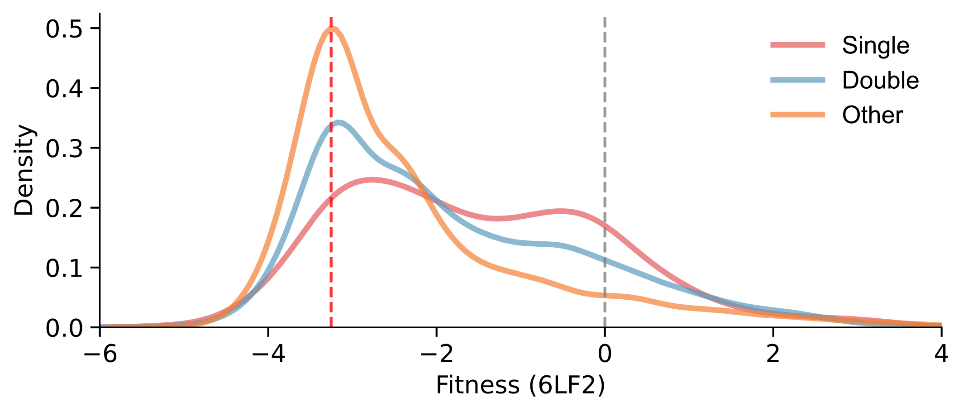

**Supplementary Figure S8.** **The distribution of fitness values for single, double and other mutants of SeviL (6LF2).** Vertical dashed lines in grey and red indicate the fitness of the wild-type variants and the median fitness of Stop codon mutations, respectively.

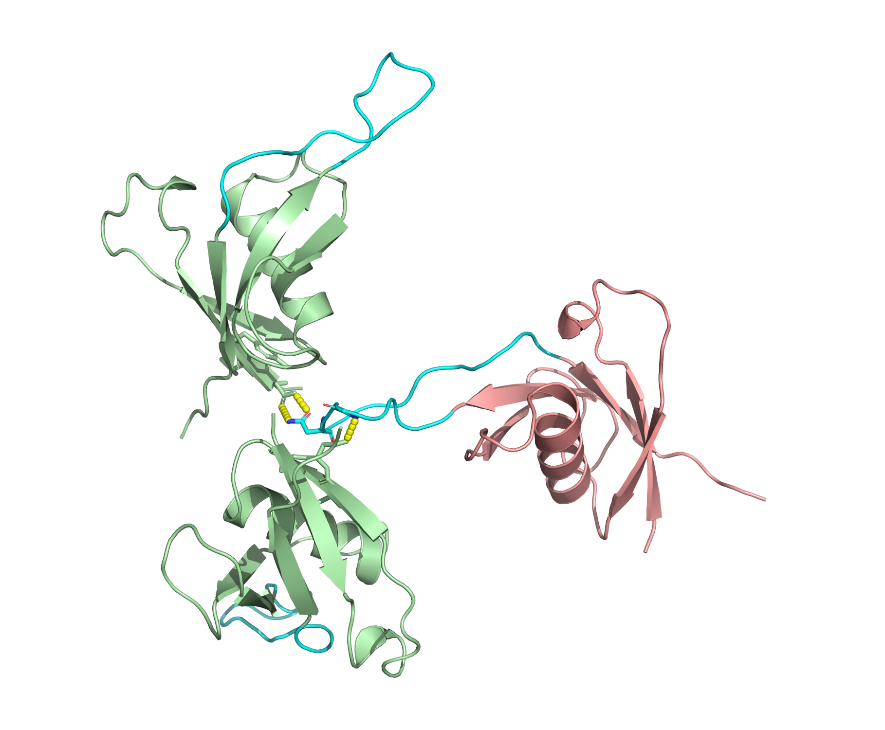

**Supplementary Figure S9.** **The crystal packing structure of 7CHQ.** The conformation of the long loop (cyan) in the β-hairpin is affected by its symmetry-related molecule (green) in the crystal packing environment. The surrounding symmetry molecules formed hydrogen bonds with the long flexible loop as shown in the figure.

**Supplementary Table S1.** Statistics of the least abundant variants before and after selections under different experimental conditions.

|  | Sequence number of the least abundant variant | | | |
| --- | --- | --- | --- | --- |
| # Exp. Conditions (μg/ml – 12 h) | Pin1 | Villin | BBL | hYAP65 |
| 25-1 Before selection | 160 | 13 | 106 | 16 |
| After selection | 238 | 43 | 277 | 13 |
| 25-2 Before selection | 160 | 13 | 106 | 16 |
| After selection | 181 | 39 | 121 | 22 |
| 25-3 Before selection | 160 | 13 | 106 | 16 |
| After selection | 175 | 44 | 127 | 7 |
| 100-1 Before selection | 160 | 13 | 106 | 16 |
| After selection | 164 | 34 | 158 | 25 |
| 100-2 Before selection | 160 | 13 | 106 | 16 |
| After selection | 273 | 61 | 194 | 23 |
| 100-3 Before selection | 160 | 13 | 106 | 16 |
| After selection | 179 | 24 | 135 | 27 |
| 400-1 Before selection | 160 | 13 | 106 | 16 |
| After selection | 156 | 20 | 107 | 25 |
| 400-2 Before selection | 160 | 13 | 106 | 16 |
| After selection | 169 | 23 | 120 | 28 |
| 400-3 Before selection | 160 | 13 | 106 | 16 |
| After selection | 168 | 27 | 131 | 21 |

**Supplementary Table S2.** Statistics of sequence variants of 6LYD before and after selections under different experimental conditions.

| # Exp. Conditions (μg/ml – 12 h) | # mutations | # sequences | Single | Double | >2 mutations |
| --- | --- | --- | --- | --- | --- |
| 25-1 Before selection | 1.80 | 2585 | 526 | 2057 | 1 |
| After selection | 1.63 | 263 | 98 | 165 | 0 |
| 25-2 Before selection | 1.79 | 2494 | 526 | 1966 | 1 |
| After selection | 1.56 | 179 | 79 | 100 | 0 |
| 25-3 Before selection | 1.78 | 2371 | 522 | 1847 | 1 |
| After selection | 1.53 | 159 | 75 | 84 | 0 |
| 100-1 Before selection | 1.80 | 2583 | 527 | 2054 | 1 |
| After selection | 1.62 | 277 | 105 | 172 | 0 |
| 100-2 Before selection | 1.79 | 2503 | 526 | 1975 | 1 |
| After selection | 1.58 | 196 | 83 | 113 | 0 |
| 100-3 Before selection | 1.78 | 2345 | 523 | 1820 | 1 |
| After selection | 1.60 | 184 | 74 | 110 | 0 |
| 400-1 Before selection | 1.80 | 2600 | 527 | 2071 | 1 |
| After selection | 1.67 | 348 | 114 | 234 | 0 |
| 400-2 Before selection | 1.79 | 2518 | 523 | 1993 | 1 |
| After selection | 1.68 | 275 | 88 | 187 | 0 |
| 400-3 Before selection | 1.78 | 2439 | 525 | 1912 | 1 |
| After selection | 1.65 | 237 | 83 | 154 | 0 |

**Supplementary Table S3.** Statistics of sequence variants of three proteins before and after selections at a given experimental condition.

| PDB ID | Structural Fold | L | # mutations | # sequences | Single | Double | >2 mutations | # Exp. Conditions |
| --- | --- | --- | --- | --- | --- | --- | --- | --- |
| 7F0K | 2-helix bundle | 92 | 3.30 | 39845 | 607 | 11675 | 27562 | 10-3^a^ |
| After selection: | | | 3.11 | 8192 | 178 | 2763 | 5251 |  |
| 6TN7 | 4-helix bundle | 97 | 1.98 | 7300 | 671 | 6128 | 500 | 25-2 |
| After selection: | | | 1.94 | 1189 | 139 | 979 | 71 |  |
| 7CHQ | Mostly sheets | 84 | 3.03 | 28012 | 600 | 11066 | 16345 | 25-2 |
| After selection: | | | 2.48 | 3101 | 165 | 1791 | 1145 |  |

^a^25 μg/ml TMP for 2 x 12h’s selection was not employed for 7F0K due to appearance of precipitation. 10 μg/ml TMP for 3 x 12h’s selection was utilized instead.

**Supplementary Table S4.** Statistics of sequence variants of 6LF2 before and after selections at a given experimental condition.

| PDB ID | L | # mutations | # sequences | Single | Double | >2 mutations | # Exp. Conditions |
| --- | --- | --- | --- | --- | --- | --- | --- |
| 6LF2 | 129 | 2.90 | 10586 | 780 | 4214 | 5591 | 10-2 ^a^ |
| After selection: | | 2.51 | 1317 | 133 | 638 | 546 |  |

^a^25 μg/ml TMP for 2 x 12h’s selection was not employed for 6LF2 due to appearance of precipitation. 10 μg/ml TMP for 2 x 12h’s selection was utilized instead.

**Supplementary Table S5.** DNA sequences used in this study.

| Name | DNA sequence (5’-3’) | Note |
| --- | --- | --- |
| mF1,2 | ATGGTGCGTCCGCTGAATTGTATTGTTGCAGTTAGTCAGAATATGGGTATTGGTAAAAATGGCGATCTGCCGTGGCCGCCGCTGCGTAATGAAAGCAAATATTTTCAGCGTATGACCACCACCAGTAGCGTTGAAGGCAAACAGAATCTGGTTATTATGGGTCGCAAAACCTGGTTTAGTATTCCGGAAAAGAATCGCCCGCTGAAAGATCGCATTAATATTGTGCTGAGTCGCGAACTGAAAGAACCGCCGCGTGGTGCCCATTTTCTGGCAAAAAGCCTGGATGATGCCCTGCGTCTGATTGAACAGCCGGAACT | Mouse DHFR (N terminal fragment) |
| mF3 | GCAAGCAAAGTGGATATGGTTTGGATTGTTGGTGGCAGTAGCGTTTATCAGGAAGCAATGAATCAGCCGGGCCATCTGCGCCTGTTTGTGACCCGCATTATGCAGGAATTTGAAAGCGATACCTTTTTCCCGGAAATTGATCTGGGTAAATATAAACTGCTGCCGGAATATCCGGGTGTGCTGAGCGAAGTTCAGGAAGAAAAAGGCATTAAGTATAAATTCGAGGTGTATGAAAAGAAGGAT | Mouse DHFR (C terminal fragment) |
| 6LYD-WT | ACCAACCTGTCTGACATCATTGAAAAAGAAACTGGCAAACAGCTGGTGATTCAGGAAAGCATCCTGATGCTGCCGGAAGAAGTTGAAGAAGTAATCGGCAACAAACCAGAGAGCGACATTCTGGTACACACCGCTTACGATGAATCTACCGACGAAAACGTGATGCTGCTGACCAGCGACGCACCGGAATACAAACCTTGGGCTCTGGTGATTCAGGATTCCAACGGTGAAAACAAAATTAAAATGCTG | DNA Template |
| 6TN7-WT | GGTGCAATGGGTACTCTGTCTCGTGAAGCTATTCAGCGTGAGCTGGATCTGCCGCAGAAACAGGGTGAACCTCTGGACCAGTTCCTGTGGCGTAAACGTGACCTGTACCAGACTCTGTACGTCGATGCGGATGAAGAAGAAATTATCCAGTACGTTGTCGGCACCCTGCAGCCGAAACTGAAGCGTTTTCTGCGTCACCCGCTGCCAAAAACCCTGGAACAGCTGATCCAGCGTGGCATGGAAGTTCAGGATGATCTGGAACAAGCAGCAGAACCGGCTGGTCCGCATCTG | DNA Template |
| 6LF2-WT | ATGAGCAGTGTTACCATTGGTAAATGTTATATTCAGAACCGCGAAAATGGCGGTCGTGCATTTTATAATCTGGGTCGCAAAGATCTGGGTATTTTTACCGGCAAAATGTATGATGATCAGATTTGGAGCTTTCAGAAAAGTGATACCCCGGGTTATTATACCATTGGCCGCGAAAGTAAATTTCTGCAGTATAATGGTGAACAGGTTATTATGAGTGATATTGAACAGGATACCACCCTGTGGAGTCTGGAAGAAGTGCCGGAAGATAAAGGCTTTTATCGCCTGCTGAATAAGGTTCATAAAGCCTATCTGGATTATAATGGCGGTGACCTGGTGGCAAATAAGCATCAGACCGAAAGCGAAAAATGGATTCTGTTTAAAGCATAT | DNA Template |
| 7CHQ-WT | ATGAACACCTATCTGATTGATCCGCGCAAAAATAATGATAATAGCGGTGAACGTTTCACCGTGGATGCCGTTGATATTACCGCAGCAGCCAAAAGTGCAGCCCAGCAGATTCTGGGCGAAGAATTTGAAGGTCTGGTGTATCGTGAAACCGGCGAAAGCAATGGCAGCGGCATGTTTCAGGCCTATCATCATCTGCATGGTACAAATCGCACCGAAACCACCGTGGGTTATCCGTTTCATGTGATGGAACTG | DNA Template |
| Evo-FP | TGGCGGCGGTAGTGGTACC | EP-PCR  Primer |
| Evo-RP | ACCGCCACCGCCGGATCC | EP-PCR  Primer |
| P5-U085-RD1-FP | AATGATACGGCGACCACCGAGATCTACACcggaactgACACTCTTTCCCTACACGACGCTCTTCCGATCTGGCGGCGGTAGTGGtaCC | Primer for HTS samples preparation |
| P7-U085-RD2-RP | CAAGCAGAAGACGGCATACGAGATtataacctGTGACTGGAGTTCAGACGTGTGCTCTTCCGATCtACCGCCACCGCCGGATCC | Primer for HTS samples preparation |
| P5-U086-RD1-FP | AATGATACGGCGACCACCGAGATCTACACtaaggtcaACACTCTTTCCCTACACGACGCTCTTCCGATCTGGCGGCGGTAGTGGtaCC | Primer for HTS samples preparation |
| P7-U086-RD2-RP | CAAGCAGAAGACGGCATACGAGATcgcggttcGTGACTGGAGTTCAGACGTGTGCTCTTCCGATCtACCGCCACCGCCGGATCC | Primer for HTS samples preparation |
| P5-U087-RD1-FP | AATGATACGGCGACCACCGAGATCTACACttgcctagACACTCTTTCCCTACACGACGCTCTTCCGATCTGGCGGCGGTAGTGGtaCC | Primer for HTS samples preparation |
| P7-U087-RD2-RP | CAAGCAGAAGACGGCATACGAGATttggtgagGTGACTGGAGTTCAGACGTGTGCTCTTCCGATCtACCGCCACCGCCGGATCC | Primer for HTS samples preparation |
| P5-U088-RD1-FP | AATGATACGGCGACCACCGAGATCTACACccattcgaACACTCTTTCCCTACACGACGCTCTTCCGATCTGGCGGCGGTAGTGGtaCC | Primer for HTS samples preparation |
| P7-U088-RD2-RP | CAAGCAGAAGACGGCATACGAGATccaacagaGTGACTGGAGTTCAGACGTGTGCTCTTCCGATCtACCGCCACCGCCGGATCC | Primer for HTS samples preparation |
| 108-oligo pool | gatggtgcgactagaggtaccATGGCCGATGAGGAAAAAGCGCCGCCGGGATGGGAAAAACGTATGTCACGTTCTTCTGGCCGCGTGTATTACTTTAACCATATTACCAATGCAAGCCAGTGGGAACGCCCAAGTGGTGGTTCGAATGGCAGCGGGAACGGCAGCTCCGGTggatccgaacaaaagcttatttctga  gatggtgcgactagaggtaccATGGCGGATGAAGAAAAAATTCCGCCGGGCTGGGAAAAACGTATGTCTCGCAGCAGTGGTCGTGTGTATTACTTTAATCATATCACCAACGCCAGCCAGTGGGAGCGCCCATCTGGCGGGAGCAATGGCTCGGGAAACGGTTCCTCAGGTggatccgaacaaaagcttatttctga  gatggtgcgactagaggtaccATGGCGGATGAAGAGAAAGTGCCGCCGGGCTGGGAAAAACGTATGTCACGTAGCTCGGGGCGCGTTTATTACTTTAATCATATTACCAACGCCTCTCAGTGGGAACGCCCATCCGGTGGTAGCAACGGCAGCGGAAATGGTTCTAGTGGCggatccgaacaaaagcttatttctga  gatggtgcgactagaggtaccATGGCCGATGAAGAAAAACTGGCGCCAGGTTGGGAAAAACGCATGTCGCGTTCCAGCGGCCGCGTGTACTATTTTAACCATATTACCAATGCATCACAGTGGGAGCGTCCGTCTGGAGGGTCTAATGGTAGCGGCAACGGTAGCAGTGGCggatccgaacaaaagcttatttctga  gatggtgcgactagaggtaccATGGCGGATGAAGAAAAACTGGGTCCAGGTTGGGAAAAACGTATGAGCCGCTCCAGCGGCCGCGTGTACTATTTTAATCATATTACCAACGCCTCGCAGTGGGAGCGTCCGAGCGGCGGATCAAACGGGAGTGGCAATGGTTCTTCTGGCggatccgaacaaaagcttatttctga  gatggtgcgactagaggtaccATGGCGGATGAAGAAAAACTGCCAGCAGGTTGGGAAAAACGTATGAGCCGCAGTTCTGGTCGCGTGTATTACTTTAATCATATTACCAACGCCAGCCAGTGGGAGCGTCCGTCCGGCGGTTCGAACGGCTCTGGCAATGGAAGCTCAGGGggatccgaacaaaagcttatttctga  gatggtgcgactagaggtaccATGGCCGATGAAGAAAAACTGCCGGGGGGCTGGGAAAAACGCATGAGTCGTTCCTCTGGTCGTGTGTATTACTTTAATCATATTACCAACGCGAGCCAGTGGGAGCGCCCATCAGGCGGAAGCAACGGCTCTGGTAATGGTTCGAGCGGCggatccgaacaaaagcttatttctga  gatggtgcgactagaggtaccATGGCCGATGAAGAGAAACTGCCACCGGCATGGGAAAAACGTATGAGCCGTTCGAGCGGCCGCGTGTACTATTTTAACCATATTACCAACGCGTCCCAGTGGGAACGCCCGAGTGGCGGATCTAATGGGTCTGGTAATGGCTCAAGCGGTggatccgaacaaaagcttatttctga  gatggtgcgactagaggtaccATGGCGGATGAAGAAAAACTGCCGCCGGGAGCCGAGAAACGCATGAGTCGCTCCTCAGGCCGTGTGTATTACTTTAACCATATTACCAACGCAAGCCAGTGGGAACGTCCAAGCGGCGGTTCTAATGGTTCTGGGAATGGTAGCTCGGGCggatccgaacaaaagcttatttctga  gatggtgcgactagaggtaccATGGCGGATGAAGAAAAATTACCGCCAGGCCTGGAGAAACGTATGTCTCGCAGCTCGGGCCGCGTGTATTACTTTAACCATATTACCAATGCCAGTCAGTGGGAACGTCCGAGCGGTGGCTCTAACGGATCCGGTAATGGTTCAAGCGGGggatccgaacaaaagcttatttctga  gatggtgcgactagaggtaccATGGCGGATGAGGAAAAACTGCCGCCGGGTTTTGAAAAACGCATGAGCCGTAGCTCGGGGCGCGTGTACTATTTCAACCATATTACCAATGCCAGCCAGTGGGAACGTCCATCTGGCGGTTCAAACGGCAGTGGCAATGGTTCTTCCGGAggatccgaacaaaagcttatttctga  gatggtgcgactagaggtaccATGGCTGATGAAGAGAAACTGCCGCCAGGCTGGGCCAAACGTATGTCCCGTTCGAGCGGTCGCGTGTATTACTTTAACCATATTACCAATGCGAGCCAGTGGGAACGCCCGTCTGGTGGCTCTAACGGTAGTGGGAATGGCTCAAGCGGAggatccgaacaaaagcttatttctga  gatggtgcgactagaggtaccATGGCGGATGAAGAGAAACTGCCGCCAGGTTGGGAAGCACGCATGAGTCGCTCTAGCGGGCGTGTGTACTATTTTAACCATATTACCAATGCCAGCCAGTGGGAACGTCCGTCGGGCGGCTCTAACGGATCAGGTAATGGTTCCAGCGGCggatccgaacaaaagcttatttctga  gatggtgcgactagaggtaccATGGCGGATGAGGAAAAACTGCCACCGGGCTGGGAAAAAGCCATGTCTCGTAGCAGCGGTCGGGTGTATTACTTTAACCATATTACCAACGCAAGCCAGTGGGAACGCCCGTCTGGCGGTTCGAATGGGTCCGGTAATGGCTCAAGTGGAggatccgaacaaaagcttatttctga  gatggtgcgactagaggtaccATGGCCGATGAAGAGAAACTGCCGCCAGGCTGGGAAAAACGCGCGAGCCGTAGCAGCGGTCGCGTGTACTATTTTAACCATATTACCAACGCATCTCAGTGGGAACGTCCGTCTGGCGGTTCAAATGGTAGTGGAAATGGCTCGTCCGGGggatccgaacaaaagcttatttctga  gatggtgcgactagaggtaccATGGCTGATGAAGAGAAACTGCCGCCGGGTTGGGAAAAACGCATGGCGCGCAGCTCGGGTCGTGTGTATTACTTTAATCATATTACCAATGCCTCTCAGTGGGAACGTCCAAGTGGCGGATCAAACGGCAGCGGGAACGGCTCCAGCGGTggatccgaacaaaagcttatttctga  gatggtgcgactagaggtaccATGGCCGATGAAGAGAAACTGCCGCCAGGCTGGGAAAAACGCATGGGACGTAGCAGCGGCCGCGTGTACTATTTTAATCATATTACCAATGCGAGCCAGTGGGAACGTCCGTCAGGCGGTAGTAACGGTTCGGGCAACGGTTCCTCTGGGggatccgaacaaaagcttatttctga  gatggtgcgactagaggtaccATGGCTGATGAAGAGAAACTGCCGCCGGGATGGGAAAAACGGATGTCGGCCAGCTCCGGTCGCGTGTACTATTTTAATCATATTACCAACGCGTCTCAGTGGGAACGTCCAAGCGGTGGCAGTAACGGGTCAGGTAATGGCTCTAGCGGCggatccgaacaaaagcttatttctga  gatggtgcgactagaggtaccATGGCCGATGAAGAAAAACTGCCACCGGGCTGGGAAAAACGGATGTCTGGCAGTTCGGGTCGCGTGTATTACTTTAATCATATTACCAACGCGTCTCAGTGGGAGCGTCCGAGCGGAGGCTCCAACGGCTCAGGGAATGGTAGCAGCGGTggatccgaacaaaagcttatttctga  gatggtgcgactagaggtaccATGGCAGATGAGGAAAAACTGCCGCCGGGCTGGGAAAAACGCATGTCTCGCGCCAGTGGACGTGTGTACTATTTTAACCATATTACCAACGCGAGCCAGTGGGAACGTCCAAGCGGGGGTTCCAATGGCTCAGGCAATGGTTCGAGCGGTggatccgaacaaaagcttatttctga  gatggtgcgactagaggtaccATGGCGGATGAAGAAAAACTGCCGCCGGGCTGGGAGAAACGTATGTCGCGTGGTAGCGGCCGCGTGTACTATTTTAATCATATTACCAACGCCTCACAGTGGGAACGCCCAAGCGGCGGTTCCAACGGCAGTGGGAATGGAAGCTCTGGTggatccgaacaaaagcttatttctga  gatggtgcgactagaggtaccATGGCCGATGAAGAAAAACTGCCACCGGGATGGGAAAAACGCATGTCCCGTTCTGCAGGCCGCGTGTATTACTTTAATCATATTACCAACGCGAGCCAGTGGGAGCGTCCGAGCGGCGGGTCAAATGGCTCGGGTAACGGTAGCAGTGGTggatccgaacaaaagcttatttctga  gatggtgcgactagaggtaccATGGCCGATGAAGAGAAACTGCCGCCAGGTTGGGAAAAACGTATGTCACGCAGTGGCGGGCGCGTGTACTATTTTAACCATATTACCAACGCGTCCCAGTGGGAACGTCCGTCGGGAGGTAGCAATGGCAGCGGCAATGGTTCTAGCGGCggatccgaacaaaagcttatttctga  gatggtgcgactagaggtaccATGGCCGATGAAGAAAAACTGCCGCCGGGTTGGGAGAAACGCATGTCGCGCTCTAGCGCACGTGTGTACTATTTTAATCATATTACCAACGCGAGTCAGTGGGAACGTCCAAGCGGAGGCTCCAACGGGTCTGGCAATGGCTCAAGCGGTggatccgaacaaaagcttatttctga  gatggtgcgactagaggtaccATGGCTGATGAAGAGAAACTGCCGCCAGGTTGGGAAAAACGCATGTCACGTAGCTCCGGAGCGGTGTACTATTTTAACCATATTACCAATGCCTCTCAGTGGGAACGGCCGAGTGGCGGGAGCAACGGTTCGGGTAATGGCAGCTCTGGCggatccgaacaaaagcttatttctga  gatggtgcgactagaggtaccATGGCGGATGAAGAAAAACTGCCGCCGGGTTGGGAGAAACGCATGAGCCGGAGCAGCGGGGGTGTGTACTATTTTAACCATATTACCAATGCCTCACAGTGGGAACGTCCATCTGGCGGCAGTAACGGCTCTGGCAATGGATCGTCCGGTggatccgaacaaaagcttatttctga  gatggtgcgactagaggtaccATGGCGGATGAGGAAAAACTGCCACCGGGATGGGAAAAACGCATGAGCCGTTCTTCTGGCCGTGCATACTATTTTAACCATATTACCAATGCCAGTCAGTGGGAACGCCCGTCCGGCGGTAGCAACGGGTCGGGCAATGGTTCAAGCGGTggatccgaacaaaagcttatttctga  gatggtgcgactagaggtaccATGGCGGATGAGGAAAAACTGCCACCGGGTTGGGAAAAACGCATGTCTCGTAGCAGTGGCCGTGTGGCCTATTTTAACCATATTACCAATGCATCCCAGTGGGAACGCCCGAGCGGTGGCAGCAACGGATCAGGCAATGGGTCGTCTGGTggatccgaacaaaagcttatttctga  gatggtgcgactagaggtaccATGGCCGATGAAGAAAAACTGCCGCCAGGTTGGGAGAAACGTATGTCTCGCTCGTCTGGCCGCGTGTTATATTTTAATCATATTACCAACGCGTCACAGTGGGAACGTCCGTCCGGCGGTAGCAACGGGAGTGGTAATGGAAGCAGCGGCggatccgaacaaaagcttatttctga  gatggtgcgactagaggtaccATGGCCGATGAAGAAAAACTGCCGCCGGGTTGGGAGAAACGTATGTCCCGCAGCTCTGGCCGTGTGTTCTATTTTAATCATATTACCAACGCGAGTCAGTGGGAACGCCCATCGGGCGGGAGCAACGGAAGCGGTAATGGCTCATCTGGTggatccgaacaaaagcttatttctga  gatggtgcgactagaggtaccATGGCCGATGAAGAAAAACTGCCGCCAGGCTGGGAAAAACGTATGTCGCGCAGCAGCGGTCGCGTGTATTTATTTAATCATATTACCAACGCGTCACAGTGGGAGCGTCCGTCTGGCGGGTCCAACGGTAGTGGAAATGGCAGCTCTGGTggatccgaacaaaagcttatttctga  gatggtgcgactagaggtaccATGGCCGATGAAGAAAAACTGCCACCGGGATGGGAGAAACGCATGAGTCGTTCCTCGGGTCGCGTGTATTTTTTCAATCATATTACCAACGCGTCTCAGTGGGAACGTCCGTCTGGGGGCAGCAACGGCTCAGGTAATGGCAGCAGCGGTggatccgaacaaaagcttatttctga  gatggtgcgactagaggtaccATGGCGGATGAAGAAAAACTGCCGCCGGGTTGGGAGAAACGTATGAGCCGCTCTTCCGGTCGTGTGTATTGGTTTAACCATATTACCAACGCCTCACAGTGGGAACGCCCAAGTGGCGGTTCGAATGGCTCTGGCAATGGAAGCAGCGGGggatccgaacaaaagcttatttctga  gatggtgcgactagaggtaccATGGCCGATGAAGAAAAACTGCCGCCAGGTTGGGAGAAACGTATGAGCCGCAGCTCTGGTCGCGTGTATTACGCGAATCATATTACCAATGCAAGTCAGTGGGAACGTCCGTCAGGCGGTTCGAACGGCAGCGGAAACGGGTCCTCTGGCggatccgaacaaaagcttatttctga  gatggtgcgactagaggtaccATGGCCGATGAGGAAAAACTGCCGCCGGGCTGGGAAAAACGTATGAGTCGCTCAAGCGGTCGTGTGTATTACTTAAATCATATTACCAACGCGAGCCAGTGGGAACGCCCATCGGGAGGTTCCAACGGTTCTGGCAATGGCTCTAGCGGGggatccgaacaaaagcttatttctga  gatggtgcgactagaggtaccATGGCCGATGAGGAAAAACTGCCGCCGGGGTGGGAAAAACGTATGAGCCGTAGCTCTGGTCGCGTGTATTACTATAACCATATTACCAACGCGAGCCAGTGGGAACGCCCATCCGGCGGCTCAAATGGCTCTGGAAATGGTTCGAGTGGTggatccgaacaaaagcttatttctga  gatggtgcgactagaggtaccATGGCTGATGAAGAGAAACTGCCACCGGGTTGGGAAAAACGCATGTCTCGCTCCAGCGGCCGTGTGTATTACTTTGCGCATATTACCAACGCCTCGCAGTGGGAACGTCCGTCTGGAGGGAGCAACGGCTCAGGTAATGGTAGCAGTGGCggatccgaacaaaagcttatttctga  gatggtgcgactagaggtaccATGGCGGATGAAGAAAAACTGCCGCCAGGTTGGGAGAAACGCATGAGCCGTAGCAGCGGTCGCGTGTACTATTTTTTACATATTACCAACGCCTCGCAGTGGGAACGTCCGTCTGGCGGCTCCAATGGCTCTGGGAACGGTTCAAGTGGAggatccgaacaaaagcttatttctga  gatggtgcgactagaggtaccATGGCGGACGAAGAAAAACTGCCGCCAGGATGGGAAAAACGTATGAGCCGCAGTAGCGGTCGCGTGTATTACTTTGATCATATTACCAACGCCTCCCAGTGGGAGCGTCCGTCTGGTGGCTCTAACGGCAGCGGGAATGGTTCATCGGGCggatccgaacaaaagcttatttctga  gatggtgcgactagaggtaccATGGCCGATGAAGAAAAACTGCCGCCAGGATGGGAAAAACGCATGTCCCGCAGCTCTGGTCGTGTGTATTACTTTAATGCGATTACCAACGCAAGCCAGTGGGAGCGTCCGTCTGGCGGCAGCAACGGTTCGGGCAATGGGTCAAGTGGTggatccgaacaaaagcttatttctga  gatggtgcgactagaggtaccATGGCCGATGAAGAAAAACTGCCACCGGGATGGGAGAAACGCATGAGCCGCTCTTCGGGCCGTGTGTACTATTTTAACGGGATTACCAATGCGTCTCAGTGGGAACGTCCGAGCGGCGGCAGCAACGGTTCAGGTAATGGTTCCAGTGGCggatccgaacaaaagcttatttctga  gatggtgcgactagaggtaccATGGCAGATGAAGAAAAACTGCCACCGGGCTGGGAGAAACGCATGAGCCGTAGCAGCGGTCGTGTGTATTACTTTAATCATGCGACCAATGCCTCCCAGTGGGAACGCCCGTCGGGTGGATCAAACGGCTCTGGCAACGGTAGTTCTGGGggatccgaacaaaagcttatttctga  gatggtgcgactagaggtaccATGGCGGATGAGGAAAAACTGCCGCCGGGTTGGGAAAAACGCATGAGCCGCTCTAGTGGCCGTGTGTACTATTTTAACCATGGCACCAACGCCTCACAGTGGGAACGTCCATCCGGAGGTTCGAATGGGAGCGGCAATGGCAGCTCTGGTggatccgaacaaaagcttatttctga  gatggtgcgactagaggtaccATGGCCGATGAAGAGAAACTGCCACCGGGCTGGGAAAAACGTATGAGTCGCTCTAGCGGACGTGTGTACTATTTTAACCATATTGCAAACGCGTCACAGTGGGAACGCCCGTCCGGTGGTTCTAATGGCAGCGGCAATGGTAGCTCGGGGggatccgaacaaaagcttatttctga  gatggtgcgactagaggtaccATGGCCGATGAAGAAAAACTGCCGCCAGGCTGGGAGAAACGTATGAGCCGTAGCTCGGGGCGCGTGTACTATTTTAATCATATTGGCAACGCGTCCCAGTGGGAACGCCCGTCTGGCGGATCAAACGGCAGCGGTAATGGTTCTAGTGGTggatccgaacaaaagcttatttctga  gatggtgcgactagaggtaccATGGCGGATGAAGAGAAACTGCCGCCGGGCTGGGAAAAACGCATGAGCCGTTCGAGCGGTCGTGTGTACTATTTTAATCATATTTCTAACGCCTCTCAGTGGGAACGCCCAAGCGGGGGTTCAAACGGTTCCGGAAATGGCAGTTCGGGCggatccgaacaaaagcttatttctga  gatggtgcgactagaggtaccATGGCGGACGAAGAAAAACTGCCGCCGGGTTGGGAAAAACGTATGAGCCGTTCGTCAGGTCGCGTGTATTACTTTAACCATATTGATAACGCCTCTCAGTGGGAGCGCCCATCCGGCGGCTCTAATGGTAGTGGGAATGGCAGCAGCGGAggatccgaacaaaagcttatttctga  gatggtgcgactagaggtaccATGGCCGATGAGGAAAAACTGCCGCCAGGCTGGGAAAAACGTATGTCCCGCAGTTCGGGTCGCGTGTACTATTTTAACCATATTACCGCGGCAAGCCAGTGGGAACGTCCGAGCGGCGGAAGCAACGGCTCAGGTAATGGTTCTTCTGGGggatccgaacaaaagcttatttctga  gatggtgcgactagaggtaccATGGCGGATGAGGAAAAACTGCCGCCGGGCTGGGAAAAACGTATGTCCCGCAGCTCTGGCCGTGTGTATTACTTTAACCATATTACCGGTGCCTCGCAGTGGGAACGCCCAAGCGGCGGCTCAAATGGTAGTGGGAACGGATCTAGCGGTggatccgaacaaaagcttatttctga  gatggtgcgactagaggtaccATGGCGGATGAGGAAAAACTGCCGCCGGGCTGGGAAAAACGCATGTCACGTTCTTCCGGGCGTGTGTACTATTTTAATCATATTACCAACGGCAGCCAGTGGGAACGCCCAAGTGGAGGCTCTAATGGTAGCGGTAACGGTTCGAGCGGCggatccgaacaaaagcttatttctga  gatggtgcgactagaggtaccATGGCGGATGAGGAAAAACTGCCACCGGGGTGGGAAAAACGTATGAGCCGCAGCTCTGGTCGTGTGTATTACTTTAATCATATTACCAACGCCGCACAGTGGGAACGCCCGAGTGGTGGTTCGAATGGCTCAGGCAACGGAAGCTCCGGCggatccgaacaaaagcttatttctga  gatggtgcgactagaggtaccATGGCGGATGAAGAAAAACTGCCGCCGGGCTGGGAGAAACGTATGTCTCGCAGCTCGGGTCGCGTGTATTACTTTAACCATATTACCAATGCCGGCCAGTGGGAACGTCCAAGCGGCGGAAGCAATGGTAGTGGGAACGGTTCATCCGGCggatccgaacaaaagcttatttctga  gatggtgcgactagaggtaccATGGCTGATGAAGAGAAACTGCCACCGGGTTGGGAAAAACGCATGTCTCGCTCCAGCGGTCGTGTGTATTACTTTAATCATATTACCAACGCCAGTGCGTGGGAACGTCCGTCAGGCGGATCGAACGGTAGCGGGAATGGCTCTAGCGGCggatccgaacaaaagcttatttctga  gatggtgcgactagaggtaccATGGCAGATGAGGAAAAACTGCCGCCAGGTTGGGAAAAACGCATGTCGCGTTCCTCAGGTCGTGTGTATTACTTTAACCATATTACCAATGCGTCTCAGGCCGAACGCCCGAGTGGCGGCTCTAACGGTAGCGGGAATGGCAGCAGCGGAggatccgaacaaaagcttatttctga  gatggtgcgactagaggtaccATGGCCGATGAGGAAAAACTGCCGCCAGGTTGGGAAAAACGCATGAGCCGTTCTAGTGGTCGTGTGTACTATTTCAACCATATTACCAATGCGTCTCAGTTTGAACGCCCGAGCGGCGGATCGAACGGCTCAGGTAATGGCTCCAGCGGGggatccgaacaaaagcttatttctga  gatggtgcgactagaggtaccATGGCCGATGAGGAAAAACTGCCGCCGGGCTGGGAAAAACGTATGAGCCGCTCAAGTGGGCGCGTGTACTATTTTAATCATATTACCAACGCATCTCAGTGGGCGCGTCCATCGGGCGGTAGCAACGGCTCCGGTAATGGTTCTAGCGGAggatccgaacaaaagcttatttctga  gatggtgcgactagaggtaccATGGCAGATGAGGAAAAACTGCCGCCAGGCTGGGAAAAACGTATGAGCCGCTCGTCTGGACGGGTGTATTACTTTAACCATATTACCAACGCGAGTCAGTGGGAAGCCCCGAGCGGTGGCAGCAATGGTTCAGGCAATGGGTCCTCTGGTggatccgaacaaaagcttatttctga  gatggtgcgactagaggtaccATGGCCGATGAAGAAAAACTGCCACCGGGATGGGAGAAACGTATGTCACGTAGTTCTGGTCGCGTGTATTACTTTAATCATATTACCAACGCAAGCCAGTGGGAACGCGCGTCTGGCGGTTCCAACGGGTCGGGCAATGGCAGCAGCGGTggatccgaacaaaagcttatttctga  gatggtgcgactagaggtaccATGGCCGATGAAGAAAAACTGCCACCGGGCTGGGAAAAACGCATGTCCCGTTCAAGCGGCCGTGTGTACTATTTTAACCATATTACCAACGCGAGCCAGTGGGAGCGCATCAGTGGAGGTAGCAATGGGTCGGGTAATGGCTCTTCTGGTggatccgaacaaaagcttatttctga  gatggtgcgactagaggtaccATGGCAGATGAGGAAAAACTGCCACCGGGATGGGAAAAACGCATGTCTCGCAGCAGTGGCCGTGTGTATTACTTTAACCATATTACCAACGCGTCCCAGTGGGAACGTCCGGCCGGTGGCTCGAATGGTAGCGGTAATGGCAGCTCAGGGggatccgaacaaaagcttatttctga  gatggtgcgactagaggtaccATGGCCGATGAAGAGAAACTGCCACCGGGCTGGGAAAAACGTATGTCTCGCAGCAGTGGTCGTGTGTACTATTTTAATCATATTACCAACGCGTCCCAGTGGGAACGCCCGGGGGGTGGCTCAAACGGAAGCGGTAATGGCTCGAGCGGCggatccgaacaaaagcttatttctga  gatggtgcgactagaggtaccTTCGAAATCCCGGATGATGTGCCACTGCCGGCCGGTTGGGAGATGGCGAAAACGAGTTCCGGCCAACGCTATTTTTACAACCATATTGATCAGACTACCACCTGGCAGGACCCTCGTAAAGGGGGTTCGAATGGCAATTCTAACAACAGCggatccgaacaaaagcttatttctga  gatggtgcgactagaggtaccTTTGAAATTCCTGATGACGTGCCATTACCGGCGGGCTGGGAGATGGCCAAAACCAGTTCCGGTCAGCGCTATTTCCTGAACCATATCACGCAAACCACTACATGGCAGGATCCGCGTAAAGGTGGGTCGAATGGCAATAGCAACAACTCTggatccgaacaaaagcttatttctga  gatggtgcgactagaggtaccTTTGAGATCCCTGATGACGTGCCACTGCCGGCCGGGTGGGAAATGGCGAAAACTTCGTCCGGCCAACGCTATTTCTTAAACCATATTACCCAGACAACGACCTGGCAGGATCCGCGTAAAGGTGGCAGTAATGGTAATAGCAACAACTCTggatccgaacaaaagcttatttctga  gatggtgcgactagaggtaccTTTGAGATTCCGGATGATGTGCCTCTGCCAGCGGGGTGGGAAATGCGTAAAACCTCGAGTGGCCAACGGTATTTCTTAAACCATATCGACCAGACTACGACCTGGCAGGATCCGCGCAAAGGCGGTTCTAATGGTAACTCCAACAATAGCggatccgaacaaaagcttatttctga  gatggtgcgactagaggtaccTTCGAGATCCCGGACGATGTGCCATTACCGGCGGGCTGGGAAATGCGCAAAACCAGCTCTGGCCAGCGTTATTTTCTGAACCATATTACTCAGACGACCACATGGCAAGATCCTCGGAAAGGGGGTAGTAACGGTAATTCCAATAACTCGggatccgaacaaaagcttatttctga  gatggtgcgactagaggtaccTTTGAGATTCCGGACGATGTGCCGCTGCCAGCGGGTTGGGAAATGCGTAAAACGTCTTCGGGCCAACGCTATTTCTACAACCATATCGATCAGACTACCACCTGGCAGGATCCTCGGAAAGGTGGGAGTAACGGCAATAGCAATAACTCCggatccgaacaaaagcttatttctga  gatggtgcgactagaggtaccTTCGAGATCCCTGATGACGTGCCGCTGCCGGCCGGCTGGGAAATGGCGAAAACGTCTAGCGGTCAGCGTTATTTTATTAACCATATTGATCAAACTACCACCTGGCAGGATCCACGCAAAGGCGGTTCGAACGGGAATAGTAATAACTCCggatccgaacaaaagcttatttctga  gatggtgcgactagaggtaccTTTGAGATTCCTGATCAGGTGCCATTACCGGCGGGGTGGGAAATGGCCAAAACTAGTTCGGGCCAGCGTTATTTCCTGAACCATATCGACCAAACGACCACCTGGCAGGATCCGCGCAAAGGTGGTTCCAATGGCAACTCTAACAATAGCggatccgaacaaaagcttatttctga  gatggtgcgactagaggtaccTTCGAGATTCCAGATGATGTGCCGTTACCTGCCGGTTGGGAAATGGCGAAAACCAGCTCCGGCCAACGCTATTTTCTGAACCATCGGGATCAGACTACGACCTGGCAGGACCCGCGTAAAGGTGGGAGTAACGGCAATTCTAATAACTCGggatccgaacaaaagcttatttctga  gatggtgcgactagaggtaccTTCGAGATCCCTGACGATGTGCCGTTACCGGCCGGTTGGGAAATGGCGAAAACTAGTTCGGGTCAGCGGTATTTTCTGAACCATATTGATCAACGCACGACCTGGCAGGATCCACGTAAAGGCGGCTCCAACGGGAATTCTAATAACAGCggatccgaacaaaagcttatttctga  gatggtgcgactagaggtaccTTTGAGATTCCGGATGACGTGCCATTACCGGCGGGTTGGGAAATGGCCAAAACCTCTAGCGGTCAACGCTATTTCCTGAATCATATCGATAAAACCACGACTTGGCAGGATCCTCGTAAGGGGGGCTCCAACGGCAACAGTAACAATTCGggatccgaacaaaagcttatttctga  gatggtgcgactagaggtaccATGCTGTCTGATGAGGACTTTATGGCAGTGTTCGGCATGACCCGCAGTGCGTTTGCCAATCTTCCGTTGTGGAAGCAACAGAACCTGAAAAAAGAAAAAGGATTATTCGGCGGGAACTCTGGTGGTAGCTCCAGCGGTTCATCGGGCAGCggatccgaacaaaagcttatttctga  gatggtgcgactagaggtaccATGCTGTCGGATGAAGACTTCAAAGCCGTGTTTGGTATGACCCGCAGCGCGTTCGCAAACTTACCGTTGTGGATGCAGCAAAATCTGAAGAAAGAGAAAGGCCTTTTTGGGGGCAACAGCGGCGGTAGCTCTTCTGGAAGTTCAGGTTCCggatccgaacaaaagcttatttctga  gatggtgcgactagaggtaccATGCTGAGCGATGAAGACTTCAAAGCAGTGTTCGGAATGACCCGCAGCGCCTTTGCGAACTTGCCGTTATGGAAACAGCAAAATCTGATGAAGGAGAAAGGCCTTTTTGGTGGGAACAGCGGTGGTTCGTCTTCTGGCTCAAGTGGCTCCggatccgaacaaaagcttatttctga  gatggtgcgactagaggtaccATGCTGTCAGACGAAGATCTGAAAGCAGTGTTTGGAATGACCCGCTCTGCCTTCGCGAATCTTCCGCTGTGGAAGCAGCAAGCTTTAATGAAAGAGAAAGGTTTGTTTGGCGGTAACAGCGGTGGGAGCTCTAGTGGCTCCTCGGGCAGCggatccgaacaaaagcttatttctga  gatggtgcgactagaggtaccATGCTGAGTGACGAGGATTTCAAAGCGGTGCTTGGCATGACCCGCTCGGCCTTTGCTAATTTACCGTTGTGGAAACAACAGGCACTGATGAAGGAAAAAGGCCTGTTTGGTGGCAACTCAGGAGGTAGCTCCTCTGGGAGCAGCGGTTCTggatccgaacaaaagcttatttctga  gatggtgcgactagaggtaccATGTTATCTGATGAAGACTTTAAGGCAGTGTTCGGAATGACCCGCAGTGCGCTGGCCAATCTGCCGCTTTGGAAACAACAGGCTCTGATGAAAGAGAAAGGCTTGTTTGGTGGCAACAGCGGCGGGTCCAGCTCTGGTTCAAGCGGTTCGggatccgaacaaaagcttatttctga  gatggtgcgactagaggtaccATGCTGTCTGATGAGGACTTGAAAGCAGTGCTGGGTATGACCCGCAGTGCTTTTGCGAACTTACCGCTGTGGAAACAACAGGCCCTTATGAAAGAAAAGGGCCTCTTCGGAGGGAATTCGGGCGGCAGCAGCTCTGGTTCCTCAGGTAGCggatccgaacaaaagcttatttctga  gatggtgcgactagaggtaccATGCTGAGCGATGAAGACCTTAAAGCCGTGTTTGGCATGACCCGCTCTGCTTTGGCGAATCTGCCGCTGTGGAAACAGCAAGCATTAATGAAAGAGAAGGGACTCTTCGGCGGTAACTCGGGGGGCAGCAGCTCCGGTTCAAGTGGTTCTggatccgaacaaaagcttatttctga  gatggtgcgactagaggtaccATGCTTTCCGATGAAGACCTCAAAGCTGTGTTAGGTATGACCCGCAGCGCACTGGCGAATTTGCCGCTGTGGAAGCAGCAAGCCCTGATGAAAGAGAAAGGCCTGTTTGGTGGCAACAGCGGAGGGTCATCTAGCGGCTCGAGTGGTTCTggatccgaacaaaagcttatttctga  gatggtgcgactagaggtaccATGCTGTCTGATGAAGACTTTAAAGCGGTGTTCGGCATGACCCGCAGCGCTTTCGCCAACCTGCCGTTACTGAAACAGCAAGCACTCATGAAAGAGAAGGGCTTGTTTGGTGGCAATTCAGGTGGGTCGTCCAGCGGTAGCAGTGGATCTggatccgaacaaaagcttatttctga  gatggtgcgactagaggtaccCAGAATAACGACGGATTGAGCCCGGCCATCCGCCGTTTACTGGCGGAATGGAACCTTGATGCCTCTGCAATTAAAGGGACCGGCGTTGGCGGTCGCCTGACGCGTGAAGATGTGGAGAAGCATCTGGCTAAAGCGGGTGGCTCGGGTAATggatccgaacaaaagcttatttctga  gatggtgcgactagaggtaccCAGAACAATGATGCGGCCTCGCCGGCGATTCGTCGCTTACTTGCCGAGTGGAACCTGGATGCATCTGCTATCAAGGGCACGGGAGTTGGGGGCCGTTTGACCCGCGAAGACGTGGAAAAACATCTGGCGAAAGCAGGCGGTAGCGGTAATggatccgaacaaaagcttatttctga  gatggtgcgactagaggtaccCAGAACAATGATGCACTGGGACCGGCGATTCGTCGCCTGCTTGCCGAATGGAACTTAGACGCGAGCGCTATCAAAGGTACCGGGGTTGGCGGCCGTCTGACGCGCGAGGATGTGGAAAAGCATTTGGCCAAAGCGGGTGGTTCTGGCAATggatccgaacaaaagcttatttctga  gatggtgcgactagaggtaccCAGAACAATGATGCCTTGTCTCCGGGCATTCGCCGCTTACTGGCCGAGTGGAATCTGGATGCTTCGGCGATCAAGGGTACCGGTGTGGGCGGCCGTCTGACGCGTGAAGACGTTGAAAAACATCTTGCAAAAGCGGGGGGTAGCGGAAACggatccgaacaaaagcttatttctga  gatggtgcgactagaggtaccCAGAACAACGACGCGCTTTCTCCGGCTGTCCGTCGTCTGTTGGCGGAATGGAATCTGGATGCAAGCGCCATTAAGGGCACCGGGGTGGGAGGCCGCCTGACGCGCGAGGATGTTGAAAAACATTTAGCGAAAGCCGGTGGCTCGGGTAATggatccgaacaaaagcttatttctga  gatggtgcgactagaggtaccCAGAATAACGATGCGCTGTCGCCGGCAGCGCGTCGTCTGCTGGCTGAGTGGAACTTGGATGCGTCTGCCATTAAGGGCACCGGTGTTGGGGGCCGCTTAACGCGCGAAGACGTGGAAAAACATCTCGCCAAAGCAGGAGGTAGCGGCAATggatccgaacaaaagcttatttctga  gatggtgcgactagaggtaccCAGAACAATGATGCGTTGAGCCCGGCCATCCGCCGTGCTCTTGCCGAATGGAACCTGGACGCATCTGCGATTAAAGGAACGGGTGTTGGCGGCCGTTTAACCCGCGAAGATGTGGAGAAACATCTGGCAAAGGCGGGGGGTTCGGGCAATggatccgaacaaaagcttatttctga  gatggtgcgactagaggtaccCAGAATAACGACGCGTTATCTCCGGCAATCCGTCGCTTGGCGGCGGAATGGAATCTTGATGCCAGCGCCATTAAAGGGACGGGTGTTGGCGGCCGCCTGACCCGTGAAGATGTGGAGAAGCATCTGGCTAAAGCAGGTGGCTCGGGAAACggatccgaacaaaagcttatttctga  gatggtgcgactagaggtaccCAGAATAACGACGCCTTGTCTCCGGCGATTCGTCGTCTGCTCGGTGAATGGAACCTGGATGCTAGTGCGATCAAAGGTACGGGCGTGGGCGGGCGCCTGACCCGCGAGGATGTTGAAAAACATTTAGCCAAGGCAGGTGGCAGCGGAAATggatccgaacaaaagcttatttctga  gatggtgcgactagaggtaccCAGAACAATGATGCCTTATCTCCGGCAATTCGTCGTCTTCTGGCTGAGTGGAATGCAGACGCGAGCGCCATCAAGGGCACGGGCGTTGGGGGACGCTTGACCCGCGAAGATGTGGAAAAACATCTGGCGAAAGCGGGTGGCTCGGGTAACggatccgaacaaaagcttatttctga  gatggtgcgactagaggtaccCAGAACAATGATGCTTTGTCTCCGGCCATCCGTCGCCTGCTGGCGGAATGGAATCTTGACGGTTCGGCCATTAAGGGAACGGGCGTTGGGGGCCGCCTGACCCGTGAGGATGTGGAAAAACATTTAGCAAAAGCGGGCGGTAGCGGTAACggatccgaacaaaagcttatttctga  gatggtgcgactagaggtaccCAGAATAACGATGCCCTCTCTCCGGCGATCCGCCGCTTACTGGCTGAGTGGAATTTGGATGCGAGCGGTATTAAAGGGACCGGCGTGGGCGGACGTCTGACGCGTGAAGACGTTGAAAAACATCTGGCCAAGGCAGGCGGTTCGGGTAACggatccgaacaaaagcttatttctga  gatggtgcgactagaggtaccCAGAATAACGACGCGTTAAGCCCGGCAATTCGCCGTCTGTTGGCGGAATGGAACCTGGATGCGTCGGCTGTTAAGGGCACCGGCGTCGGTGGCCGCCTGACGCGTGAGGATGTGGAAAAACATCTTGCCAAAGCCGGTGGGTCTGGAAATggatccgaacaaaagcttatttctga  gatggtgcgactagaggtaccCAGAACAATGACGCCCTGTCTCCGGCAATTCGCCGTCTTTTAGCGGAATGGAATCTGGATGCCTCCGCTATCAAGGGTTCGGGCGTTGGCGGCCGCCTGACCCGTGAAGATGTGGAGAAACATTTGGCGAAAGCGGGAGGTAGCGGGAACggatccgaacaaaagcttatttctga  gatggtgcgactagaggtaccCAGAATAATGATGCGTTATCGCCGGCTATCCGTCGCCTGCTGGCCGAGTGGAACCTGGATGCGAGCGCAATTAAAGGCGCAGGTGTTGGGGGACGCCTCACCCGTGAAGACGTGGAAAAACATTTGGCCAAGGCGGGTGGCTCTGGCAACggatccgaacaaaagcttatttctga  gatggtgcgactagaggtaccCAGAATAACGATGCGTTGAGCCCGGCGATTCGTCGTCTTCTGGCTGAATGGAACCTGGACGCCTCGGCGATCAAAGGGACGGGTGGTGGCGGTCGCTTAACCCGCGAAGATGTGGAGAAGCATCTGGCAAAAGCCGGCGGATCTGGCAATggatccgaacaaaagcttatttctga  gatggtgcgactagaggtaccCAAAACAACGATGCATTGAGCCCGGCGATCCGTCGTCTTCTGGCGGAATGGAATTTAGATGCTTCTGCAATTAAGGGCACCGGTGTTGGCGGCCGCGCCACGCGCGAAGACGTGGAGAAACATCTGGCGAAAGCCGGGGGATCGGGTAATggatccgaacaaaagcttatttctga  gatggtgcgactagaggtaccCAGAACAATGACGCATTGTCCCCGGCGATTCGCCGCTTACTGGCGGAGTGGAATCTGGATGCGAGCGCCATCAAAGGCACCGGAGTTGGTGGTCGTCTGTCGCGTGAAGATGTGGAAAAGCATCTTGCCAAAGCTGGCGGCTCTGGGAACggatccgaacaaaagcttatttctga  gatggtgcgactagaggtaccCAGAATAACGATGCGTTGTCGCCGGCTATCCGCCGTCTTCTGGCGGAGTGGAATCTGGACGCCAGCGCGATTAAAGGTACGGGCGTGGGCGGACGTTTAACCCGCGAAAACGTTGAAAAACATCTGGCCAAGGCAGGTGGGTCTGGCAACggatccgaacaaaagcttatttctga  gatggtgcgactagaggtaccCAGAACAATGACGCTTTGTCGCCGGCAATTCGCCGTCTGTTAGCGGAGTGGAATCTGGATGCCTCTGCAATCAAAGGAACGGGCGTGGGCGGTCGCCTCACCCGTGAAGATGCGGAAAAGCATCTGGCCAAAGCGGGGGGTAGCGGCAACggatccgaacaaaagcttatttctga  gatggtgcgactagaggtaccCAGAACAATGATGCTTTGTCTCCGGCCATCCGCCGTCTTCTGGCAGAATGGAACCTGGATGCGAGCGCAATTAAAGGAACGGGTGTGGGCGGGCGTTTAACCCGCGAGGACGTTGAAAAAGCCCTGGCGAAGGCGGGCGGCTCGGGTAATggatccgaacaaaagcttatttctga  gatggtgcgactagaggtaccCAGAATAATGACGCCCTGTCGCCGGCGATCCGCCGTCTGTTGGCCGAGTGGAACTTAGATGCGAGCGCTATTAAAGGTACGGGCGTGGGCGGTCGTCTGACCCGCGAAGATGTTGAAAAGGGTCTTGCGAAAGCAGGGGGATCTGGCAACggatccgaacaaaagcttatttctga  gatggtgcgactagaggtaccCAGAACAATGATGCATTGAGCCCGGCGATTCGCCGCCTGCTGGCCGAATGGAATTTAGACGCGTCGGCCATCAAGGGTACCGGCGTTGGAGGTCGTCTTACGCGTGAGGATGTGGAAAAACATGCAGCGAAAGCTGGCGGCTCTGGGAACggatccgaacaaaagcttatttctga  gatggtgcgactagaggtaccCAGAATAACGATGCTTTGTCTCCGGCCATCCGCCGTCTGTTAGCGGAATGGAACCTGGATGCGAGCGCCATTAAAGGCACCGGTGTGGGCGGCCGCCTTACGCGTGAAGACGTTGAGAAGCATCTGGGAAAAGCAGGTGGGTCGGGTAATggatccgaacaaaagcttatttctga  gatggtgcgactagaggtaccCAGAATAACGATGCGTGGTCTCCGGCGATTCGTCGTCTGTTGGCGGAACATAACCTTGATGCATCGGCTATCAAGGGTACGGGAGTTGGCGGTCGCCTGACCCGCGAAGACGTGGAGAAACACTTAGCCAAAGCCGGCGGCAGCGGGAATggatccgaacaaaagcttatttctga  gatggtgcgactagaggtaccCAGAATAATGATGCGTGGTCTCCGGCGATCCGTCGTCTGCTGGCCGAGCACAACTTGGACGCCTCGGCAATTAAAGGAACGGGCGTGGGCGGGCGCTTAACCCGCGAAGATGTTGAAAAACATGCAGCGAAGGCTGGCGGTAGCGGTAACggatccgaacaaaagcttatttctga | Oligo pool synthesized by IDT |
